# Systematic detection of functional proteoform groups from bottom-up proteomic datasets

**DOI:** 10.1101/2020.12.22.423928

**Authors:** Isabell Bludau, Max Frank, Christian Dörig, Yujia Cai, Moritz Heusel, George Rosenberger, Paola Picotti, Ben C. Collins, Hannes Röst, Ruedi Aebersold

## Abstract

The cellular proteome, the ensemble of proteins derived from a genome, catalyzes and controls thousands of biochemical functions that are the basis of living cells. Whereas the protein coding regions of the genome of the human and many other species are well known, the complexity and composition of proteomes largely remains to be explored. This task is challenging because mechanisms including alternative splicing and post-translational modifications generally give rise to multiple distinct, but related proteins – proteoforms – per coding gene that expand the functional capacity of a cell.

Bottom-up proteomics is a mass spectrometric method that infers the identity and quantity of proteins from the measurement of peptides derived from these proteins by proteolytic digestion. Whereas bottom-up proteomics has become the method of choice for the detection of translation products from essentially any gene, the inherent missing link between measured peptides and their parental proteins has so far precluded the systematic assessment of proteoforms and thus limited the resolution of proteome maps. Here we present a novel, data-driven strategy to assign peptides to unique functional proteoform groups based on peptide correlation patterns across large bottom-up proteomic datasets. Our strategy does not fully characterize specific proteoforms, as is achievable in top-down approaches. Rather, it clusters peptides into functional proteoform groups that are directly linked to the biological context of the study. This allows the detection of tens to hundreds of proteoform groups in an untargeted fashion from bottom-up proteomics experiments.

We applied the strategy to two types of bottom-up proteomic datasets. The first is a protein complex co-fractionation dataset where native complexes across two different cell cycle stages were resolved and analyzed. Here, our approach enabled the systematic detection and evaluation of assembly specific proteoforms at an unprecedented scale. The second is a *protein abundance vs. sample* data matrix typical for bottom-up cohort studies consisting of tissue samples from the mouse BXD genetic reference panel. In either data type the method detected state-specific proteoform groups that could be linked to distinct molecular mechanisms including proteolytic cleavage, alternative splicing and phosphorylation. We envision that the presented approach lays the foundation for a systematic assessment of proteoforms and their functional implications directly from bottom-up proteomic datasets.

## Introduction

Human cells are known to perform thousands of different biochemical functions and the central dogma of biology states that proteins that catalyze the vast majority of these functions arise from the transcription and translation of the information contained in the respective genome. The International Human Genome Sequencing Consortium reported ~ 20,000 protein-coding genes in the human genome^1^ and surprisingly, the number of protein coding genes does not scale with the complexity of functions of eukaryotic organisms^2^. These findings have led to the notion that the protein coding information of the genome is substantially diversified structurally and functionally along the axis of gene expression^3^. Specific mechanisms that catalyze this diversification include alterative splicing of transcripts, post-translational processing and modification of proteins, and the variable association of proteins in functional protein complexes. Consequently, protein coding genes frequently give rise to multiple distinct protein species – proteoforms – that have a unique primary amino acid sequence and localized post-translational modifications (PTMs)^4,5^ and that might, in turn, partition into different functional complexes or show functional differences. Currently it is estimated that the ~ 20,000 coding genes generate more than a million different proteoforms^6^ that can differ between individual cells, tissues and disease phenotypes^4,7–9^. This increase in complexity beyond the directly translated genomic sequence information hampers genotype-based phenotype inference and highlights the importance of capturing proteome diversity to increase the mechanistic understanding of biochemical processes in basic and translational research.

Over the last decades, mass spectrometry (MS) has emerged as the key technology for proteomic analyses^10,11^. The large array of mass spectrometric techniques can be grouped into two main approaches: top-down and bottom-up proteomics. In top-down workflows, samples containing intact proteins are chromatographically separated, ionized and analyzed in a mass spectrometer. Recorded spectra of both the intact and fragmented proteins determine the unique primary protein sequence and PTMs of individual proteoforms^12^. Recent top-down proteomic studies reported the identification of more than 3,000 unique proteoforms originating from up to ~ 1,000 individual genes^13,14^. Gaining deeper proteoform coverage by top-down proteomics is challenged by the limitations of current separation techniques, the MS and MS/MS analysis of large ions, and the interpretation of the resulting spectra by available analysis software^12,15^. Furthermore, whereas top-down proteomics provides unprecedented insights into proteoform diversity, the association of proteoforms with biological function remains challenging.

Bottom-up proteomics is the more widely used technique for proteome-wide studies because some of the technical challenges facing top-down proteomics are alleviated. Here, proteins are enzymatically digested into smaller peptide sequences, which are subsequently separated by liquid chromatography (LC), ionized and analyzed by tandem mass spectrometry (MS/MS). The identity and quantity of proteins in the tested sample are subsequently inferred from the peptides that are identified based on the acquired precursor and fragment ion spectra. The method is technically robust and has demonstrated the detection of translation products of the vast majority of coding genes in a number of species. However, bottom-up proteomic workflows suffer from the principle limitation that the connectivity between identified peptides and their proteins of origin is lost during the enzymatic digestion step. This necessitates an *in silico* inference step that maps measured peptide signals back to individual proteins. This is a challenging task in general^16^ and is particularly hard for resolving different proteoforms^3^. Recent advances in instrumentation, data acquisition and data analysis, especially the development of data-independent acquisition (DIA/SWATH-MS) strategies, have enabled the measurement of large bottom-up proteomic datasets at high proteome coverage, combined with consistent and accurate quantification^17–19^. Based on these developments peptide-level bottom-up proteomic data became more reliable both on the qualitative and quantitative level, as demonstrated in several of our previous studies^20,21^. Thus, useful information about the presence of individual modifications or sequence variants on the peptide level can be readily obtained. However, the possibilities to systematically assign and distinguish unique proteoforms from bottom-up proteomics datasets remains a mostly unexplored area to date.

Nevertheless, researchers in the early days of bottom-up proteomics already observed that peptides of the same protein might follow distinct quantitative patterns across a dataset and that peptide co-variation analysis can be leveraged to improve proteomic analyses on different levels. The predominant focus of previous work has been to use peptide correlation analysis for the purpose of filtering out dissimilarly behaving peptides in an effort to improve protein quantification^22,23^ or protein inference^24^. It has also been recognized that some of the determined ‘outlier’ peptides could indeed contain valuable biological information e.g. by originating from different proteoforms and previous work explored the possibility to use peptide correlation patterns for proteoform assignment^23,25–27^. In this manuscript we describe COPF, a novel strategy for **Co**rrelation based functional **P**roteo**F**orm assessment in bottom-up proteomics data that extends the concept of peptide correlation analysis towards establishing a generic workflow with the main purpose to systematically assign peptides to co-varying proteoform groups (also see Glossary in Supplementary Table 1) in different types of bottom-up proteomics datasets.

Even more demanding than the identification of a proteoform as a distinct chemical entity is the assessment of its biological context. Although some proteoforms have successfully been annotated with molecular functions and implicated phenotypic traits^7^, the systematic assessment of proteoform-specific functions remains challenging. A shift of focus from the mere enumeration of various proteoforms detected from a cell towards establishing direct links between proteoform species and their functional significance would be a major advance in the field.

Alternative proteoforms from the same gene can functionally diverge by different mechanisms. These include i) proteoform-specific association with protein complexes, ii) different biochemical function of proteoforms and iii) different functional state of proteoforms. The COPF strategy presented here was designed to directly link proteoform group detection to specific functional contexts based on bottom-up proteomics data. Key properties of applicable datasets include both high sequence coverage and quantitative accuracy as well as a sufficient number of replicates with respect to the expected effect size of abundance differences between proteoforms across the investigated biological conditions. We recently presented a novel strategy for detecting and quantifying protein complexes from co-fractionation MS datasets^21,28,29^. The method consists of native cell lysis to extract protein complexes, followed by size-exclusion chromatography (SEC) and analysis by DIA/SWATH-MS. Because of the rich biological context, high sequence coverage and accurate quantitative peptide-level information inherent to SECSWATH-MS, we argued that it provides a unique opportunity to evaluate proteoform groups that are relevant for protein complex organization. Therefore, we first demonstrate the application of COPF to a SEC-SWATH-MS dataset where native complexes across two different cell cycle stages were resolved and analyzed^28^. In this dataset, COPF determined both assembly- and cell cycle-specific proteoform groups. We further argue that the general concept of peptide covariation based proteoform assignment should also be broadly applicable to large bottom-up proteomic studies with a high degree of biological variation. As a second application, we therefore applied COPF to assign functional proteoform groups in a typical bottom-up proteomic cohort study consisting of five tissue samples from the mouse BXD genetic reference panel^30^. In this dataset, COPF could determine several tissue-specific proteoform groups.

Overall, proteoform groups detected in either dataset could be linked to distinct molecular mechanisms including proteolytic cleavage, alternative splicing and phosphorylation. Because the strategy is agnostic to different types of proteoforms we expect that the range of applications will expand in the future and that this approach, therefore, lays the foundation for the systematic assessment of proteoform groups across large bottom-up proteomic datasets and for linking these groups to biological function.

## Results

### Principle of the method and implementation

The assignment of peptides to unique proteoforms is a challenging task in bottom-up proteomic workflows, because the majority of detected peptides are frequently shared between multiple proteoforms and multiple diverging peptides cannot be uniquely assigned during protein inference. Here we propose a novel, data-driven strategy to assign peptides to unique functional proteoform groups based on peptide correlation patterns across large bottom-up proteomic datasets (**Co**rrelation based functional **P**roteo**F**orm assessment, COPF). We define a functional proteoform group as a group of peptides, derived from the same gene, that co-vary across a large multi-condition dataset. A proteoform group can, but does not have to represent a unique, specific proteoform (also see Glossary in Supplementary Table 1). The COPF strategy is based on the following considerations: i) In case only one proteoform is expressed or all proteoforms of a protein have similar characteristics across a multi-condition dataset, all sibling peptides (i.e. peptides originating from the same parental gene/protein) should display a similar quantitative profile, as schematically illustrated in Figure 1, left panel; ii) if a gene generates multiple distinct proteoforms that differ between the analyzed samples across a multi-condition dataset, sibling peptides of that protein can be separated into groups of highly correlated peptides, which we accordingly assign to distinct proteoform groups, schematically illustrated in Figure 1, right panel.

**Figure 1.**
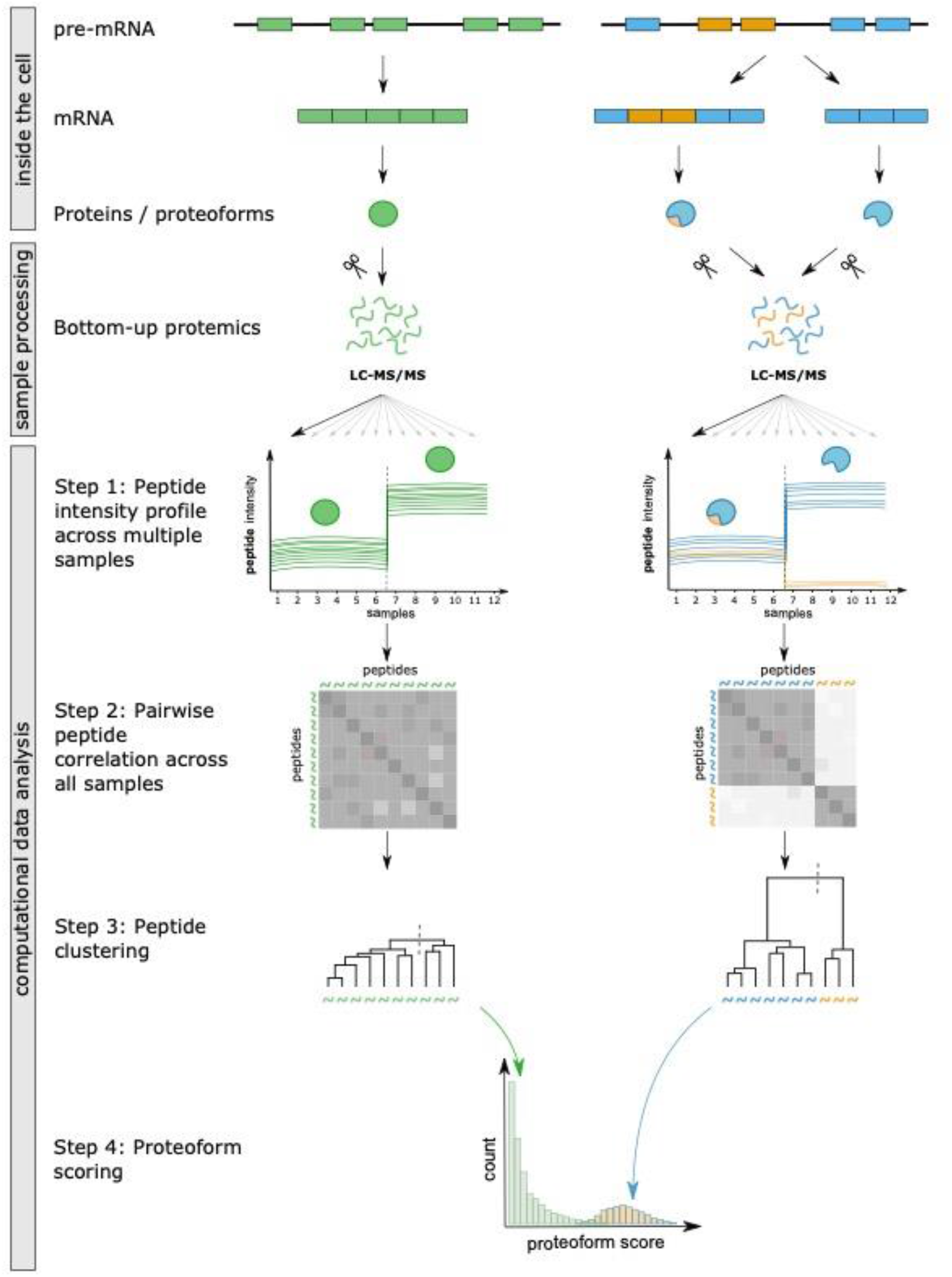
Analysis concept and workflow overview. There are multiple mechanisms by which cells can generate different proteoforms from a single gene locus. One way is via alternative splicing, which is depicted as an exemplary mechanism on the top right. The pre-mRNA contains both exons (colored boxes) and introns (lines). During alternative splicing, introns are removed and different exons can potentially be spliced in or out to generate a mature mRNA that is translated to proteins. If different splice variants are expressed in a cell, this results in multiple proteoforms with potentially different biological functions. In this example, one splice variant only contains blue exons, thus resulting in a blue proteoform. The second splice variant additionally contains two orange exons, thus generating a mixed blue and orange proteoform. During proteolytic digestion for bottom-up proteomics, proteins and proteoforms are cleaved into small peptides, a process during which the direct link between a peptide and its parental proteoform is initially lost. Our computational proteoform detection workflow is based on the concept of assessing peptide-level co-variance across larger datasets (Step 1). Here, this is illustrated by 12 samples measured across two different conditions. If there is only a single proteoform present in the dataset, all peptides should follow the same quantitative profile across all samples, here illustrated by the increase in abundance in samples of the second condition (samples 7-12, left side, green). If two different proteoforms are expressed and they are differentially regulated between conditions, only peptides of the same proteoform should follow the same trend whereas the peptides that differentiate the proteoforms show a different quantitative pattern (right side, blue peptide abundance increases versus orange peptide abundance decreases). To quantify co-variance, all pairwise peptide correlations are calculated for each protein (Step 2). The negative correlation is subsequently used for hierarchical peptide clustering into two groups (Step 3). Finally, a proteoform score is calculated based on the within-versus across-cluster correlation (Step 4). Proteins without alternative proteoforms between conditions should get a low score (green distribution), while proteins with multiple detected proteoforms between conditions should achieve a higher proteoform score (blue/orange distribution).

Conceptually, the proteoform detection workflow in COPF can be divided into four steps (Figure 1, computational data analysis). First, the intensities of peptides assigned to the same gene or protein identifier are determined from the corresponding MS signals across all measured samples. Second, all pairwise peptide correlations within a protein are calculated based on the determined intensity values across samples. Third, the peptides of a protein are subjected to hierarchical clustering, using one minus the previously calculated correlation as the distance metric. The tree is then cut into two clusters, minimally containing two peptides each (see Method section for details). Fourth, a proteoform score is calculated for each protein. The score is calculated as the mean peptide correlation across clusters minus the within-cluster correlation. A higher proteoform score thus indicates a higher within-cluster versus across-cluster correlation. The assumption of our COPF strategy is that proteins with multiple distinct proteoforms, that behave differentially across a dataset will have higher proteoform scores than proteins without differentially behaving proteoforms.

The COPF strategy is implemented and openly available as an extended version of our previously published CCprofiler R package^21,29^ at: https://github.com/CCprofiler/CCprofiler.

### Benchmark

A particular challenge for benchmarking the COPF strategy for assigning functional proteoform groups is the lack of available biological ground truth data. To evaluate COPF performance, we performed the following analyses: First, we conducted a sensitivity analysis based on an *in silico* generated benchmarking dataset. Second, we simulated the impact of different dataset properties on correct proteoform detection and grouping. Finally, we determined a set of biologically motivated criteria to evaluate the credibility of assigned proteoform groups in any given dataset.

### Benchmarking by *in silico* sensitivity analysis

As a first performance evaluation step we assessed the sensitivity of the proteoform detection workflow in COPF based on an *in silico* generated benchmarking dataset. The basis for this dataset is our previously published complex co-fractionation dataset of the HEK293 cell line. The dataset consists of native complexes isolated from a single biological condition that were separated by size-exclusion chromatography (SEC) and then analyzed by bottom-up proteomic analysis using data-independent acquisition mass spectrometry (DIA/SWATH-MS) (details on the data structure are provided in the reference and further down)^21^. We assumed that the 6299 proteins detected in this study exist as single forms and do not include any regulated proteoform groups (see below for further discussion of this assumption); we used these as a negative reference set. Exemplary peptide profiles for the NIBL1 protein are shown in Figure 2A, top panel. To obtain a positive protein reference set that COPF should classify as having multiple proteoforms, we generated an additional 3000 ‘mixed’ proteins by assigning peptides of two different parental proteins to the same pseudo mixed protein. Exemplary peptide profiles for a mixed protein based on 50% peptides of each the NADAP and the RL14 protein are shown in Figure 2A, bottom panel. The resulting set of single and mixed proteins was used to evaluate the ability of our algorithm to correctly distinguish single and mixed proteins as well as to assign peptides correctly to their corresponding parental proteins. Figure 2B shows the score distribution for the single and mixed proteins. The receiver operating characteristic (ROC) curve (Figure 2C) illustrates the dependence of true and false positive rates on the proteoform score threshold (marker color) and the percentage of mixed proteins for which all peptides were correctly assigned to their parental proteins (marker size). Overall the results of the *in silico* sensitivity analysis reveal that our hypothesis that mixture proteins can be differentiated from single proteins by means of the proteoform score holds true. At a falsepositive rate of 5.6%, 61.7% of the mixed proteins were correctly detected. For 76.8% of these mixed proteins, all peptides were completely and correctly assigned to their parental proteins. It is important to keep in mind that these are conservative estimates for the method performance for the following reasons: first, there will likely be true proteoform groups in the negative background dataset of the HEK293 sample, which are here treated as false positives. Second, some of the proteins that were grouped into one mixed protein are likely to be involved in protein interactions or having a similar elution profile by chance^29^. Peptides of such mixed proteins would therefore not be detectable as separate proteoform groups based on the correlation based COPF approach, making some of the ‘positive’ mixture proteins impossible to detect. Despite these limitations, the underlying real proteomics data used for this *in silico* benchmark has the benefit of having realistic general properties related to the number of peptides per protein as well as the measurement noise between samples. However, while we set the ratio of peptides in the mixed proteins to 50% in this analysis, this ratio might vary for actual proteoforms.

**Figure 2.**
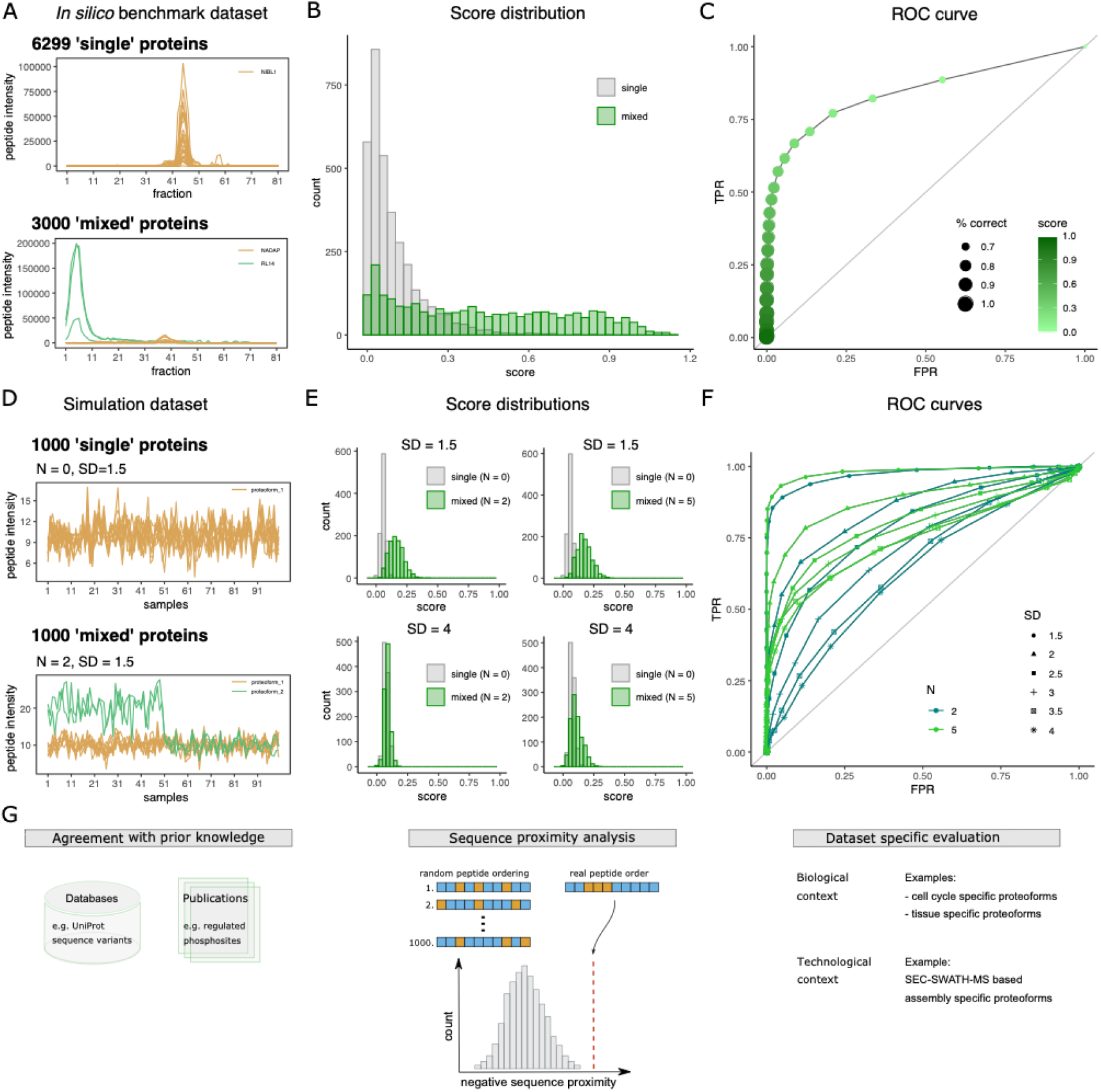
Benchmarking workflow. **(A)** Due to a lack of available ground truth proteoform data, we generated an *in silico* benchmark dataset to perform a sensitivity analysis. As a negative protein reference, we took the SEC-SWATH-MS dataset of the HEK293 proteome (details on the data structure are provided in the reference and further down)^21^. This dataset contains 6299 ‘single’ proteins. Here, peptide-level profiles of NIBL1 are shown as a representative example of a single protein i.e a protein that is detected as a single proteoform (top). To this dataset of single proteins we added 3000 pseudo ‘mixed’ proteins, each containing 50% of peptides from any two random single proteins. As an example, peptide profiles for the NADAP (orange) plus RL14 (green) mixed protein are shown. The *in silico* benchmark dataset was used to test if our workflow can correctly identify mixed proteins and assign peptides correctly to their parental proteins. **(B)** Proteoform score distribution for the single (grey) and mixed (green) proteins of the *in silico* benchmark dataset. **(C)** Receiver operator characteristic (ROC) curve for the *in silico* benchmark dataset. The marker color indicates the selected proteoform score threshold and the marker size denotes the percentage of mixed proteins for which all peptides were correctly assigned to their parental proteins. **(D)** Simulation studies were performed to evaluate the parameters that influence the success of our approach. Data for 1000 proteins was simulated by random sampling from a normal distribution across 100 samples (mean = 10, standard deviation = 1). The resulting protein profiles were used as basis to simulate 10 peptides for each protein by adding peptide-specific technical noise, sampling from a normal distribution (mean = protein intensity and standard deviation (SD) = 1.5, 2, 2.5, 3, 3.5 or 4). Exemplary simulated peptide intensity profiles for a single protein (SD = 1.5) are shown in the top panel. In addition to the 1000 single proteins, we further simulated 1000 ‘mixed’ proteins. Mixed proteins were generated by multiplying the intensity values of the first 50 samples by two for either two or five of the 10 peptides per protein (N = 2 or 5). Exemplary simulated peptide intensity profiles for a mixed protein (N = 2 and SD = 1.5) are shown in the bottom panel. **(E)** Proteoform score distributions for the single (grey) and mixed (green) proteins of the simulated datasets. **(F)** Receiver operator characteristic (ROC) curves for the simulated datasets. The color indicates the number of peptides (N) in the differentially behaving proteoform. The marker shape indicates the technical measurement noise (SD). **(G)** Credibility criteria for the detected proteoform groups in any given dataset. For the sequence proximity analysis (center), peptides are indicated as small boxes, colored by their assigned proteoform group in orange or blue.

### Benchmarking simulation study

In addition to the *in silico* sensitivity analysis, we further performed simulation studies to evaluate the impact of different dataset properties on correct proteoform detection and grouping. The simulation was performed purely based on random sampling from normal distributions without any underlying experimental data. In contrast to the *in silico* benchmark described above, this allowed us to control all parameters and to evaluate their impact on COPF performance. First, we simulated intensity profiles for 1000 ‘single’ proteins. For each protein, intensities across 100 samples were generated by random sampling from a normal distribution (mean = 10, standard deviation = 1). The resulting protein profiles were used as basis to simulate 10 peptides for each protein by adding peptide-specific technical noise, sampling from a normal distribution (mean = protein intensity and standard deviation (SD) = 1.5, 2, 2.5, 3, 3.5 or 4). Exemplary simulated peptide intensity profiles for a single protein (SD = 1.5) are shown in Figure 2D, top panel. In addition to the 1000 single proteins, we further simulated 1000 ‘mixed’ proteins. Mixed proteins were generated by multiplying the intensity values of the first 50 samples by two for either two or five of the 10 peptides per protein (N = 2 or 5). Exemplary simulated peptide intensity profiles for a mixed protein (N = 2 and SD = 1.5) are shown in Figure 2D, bottom panel. By testing different combinations of N and SD values, we could evaluate their impact on the proteoform score distribution (Figure 2E) and ROC curves (Figure 2F). The results show that 2 out of 10 peptides in a differentially behaving proteoform are sufficient to separate mixed from single proteins at low technical noise (SD = 1.5). However, increasing the measurement noise (while keeping the fold change constant) results in mostly overlapping score distributions, as can be seen in Figure 2E, bottom left (N = 2 and SD = 4). In such cases, increasing the number of peptides that distinguish the proteoforms (N = 5) improves detectability, as observed by better score separation (Figure 2E, bottom right) and an increased area under the ROC curve (Figure 2F). The results of the simulation study demonstrate the boundaries of the COPF approach. As expected, increasing the technical variation between samples at a given proteoform-specific fold change decreases the sensitivity of proteoform detection. For such cases, proteoform group detectability is better for proteoform groups with multiple distinguishing peptides as compared to proteoforms with only two distinguishing peptides.

### Benchmarking by means of auxiliary data

To finally assess and characterize the proteoform groups determined in a specific dataset, we propose evaluation criteria based on auxiliary data on three different levels: (1) agreement with prior knowledge, (2) sequence proximity analysis and (3) dataset specific criteria (Figure 2G). We demonstrate these evaluations in the two applications (below). Prior knowledge can be derived from public databases or prior publications. Here, we assessed the agreement of proteoform groups derived by the purely data driven COPF approach with annotated sequence variants in UniProt^31^. We further evaluated overlapping information with previously reported regulated phosphorylation sites (see below). For the sequence proximity analysis, we assume that many proteoforms differ by extended sequence stretches, e.g. when generated by alternative splicing or proteolytic cleavage. The sequence proximity analysis evaluates if peptides assigned to the same proteoform group are in closer relative sequence proximity than expected for random peptide grouping. Figure 2G (center) schematically illustrates the sequence proximity analysis strategy (for details also see Methods section). Finally, a key promise of the COPF approach is that the derived proteoforms can directly be linked to functional or phenotypic characteristics of the studied dataset. The reason for this is that proteoform groups are only detectable by COPF if they show different behavior across the studied dataset. Detected proteoform groups can therefore be assessed for their agreement with known underlying technical or biological structures of the given dataset, e.g. different biological conditions.

The three presented evaluation criteria can be applied to each dataset analyzed by COPF to further prioritize proteoform candidates to find promising candidates for follow-up analyses.

### Identification of cell cycle- and assembly-specific proteoforms in a SEC-SWATH-MS dataset

We applied the COPF strategy to our previously published native complex co-fractionation dataset of Hela CCL2 cells synchronized in interphase and mitosis^28^ to identify cell-cycle and assembly-specific proteoforms. In this case the dataset consists of native complexes isolated from cells in two cell cycle states, separated by size-exclusion chromatography (SEC), and then analyzed by bottom-up proteomic analysis using data-independent acquisition mass spectrometry (DIA/SWATH-MS)^21,29^. The workflow with individual steps is schematically illustrated in Figure 3A. First, cells are lysed under close to native conditions to keep protein complexes intact. Second, the protein complex mixture is separated by SEC into 65 fractions. Third, each sampled fraction is separately processed for bottom-up proteomic measurements by SWATH-MS^17^ followed by peptidecentric analysis^32–34^o. In a fourth step, peptide elution profiles across the SEC fractions are evaluated to infer protein complex assemblies by CCprofiler as described before^21,29^ and additionally by the COPF method to identify proteoforms differentially associating with complexes within or across cell cycle states.

**Figure 3.**
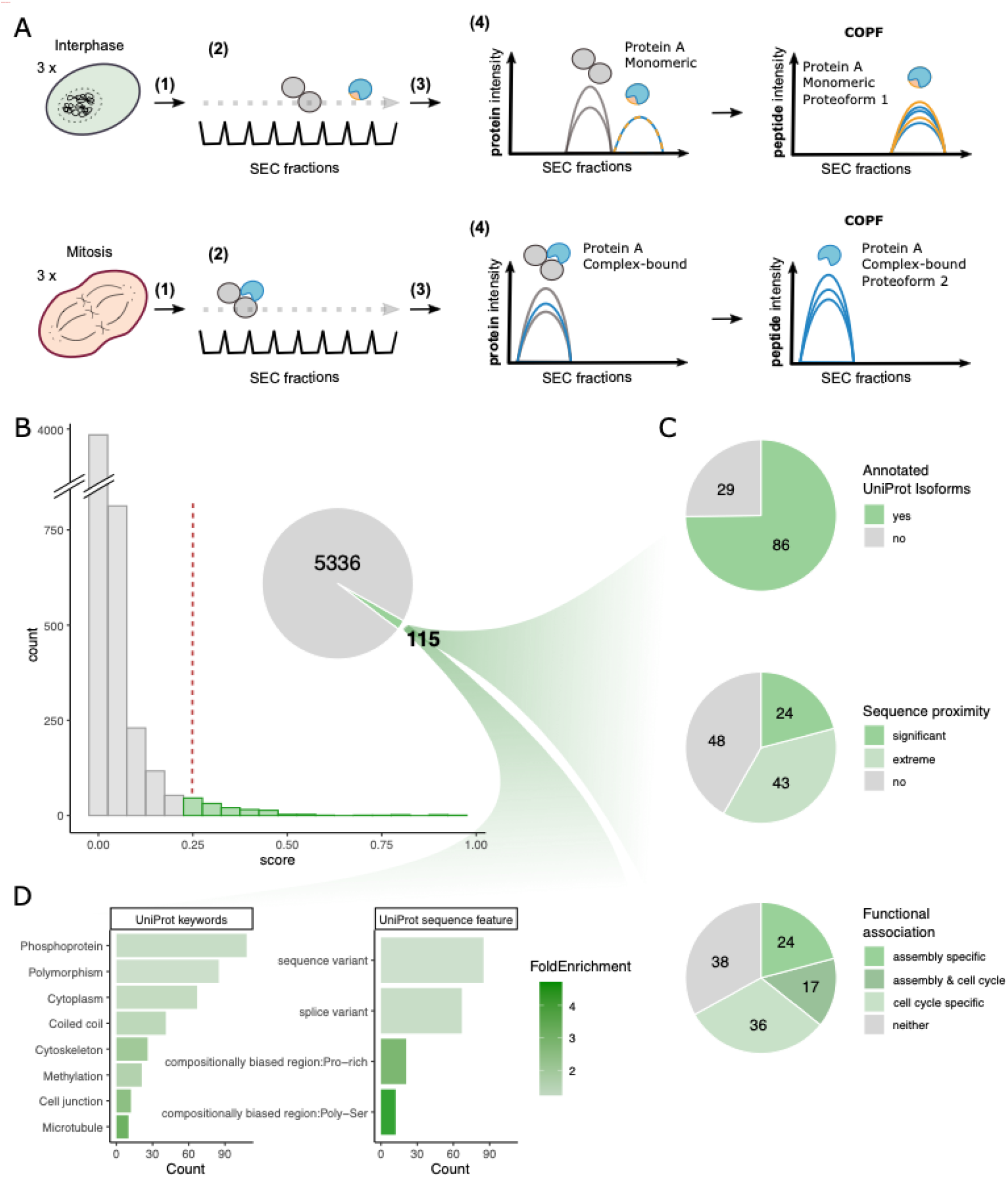
Global insights into cell cycle- and assembly-specific proteoforms in SEC-SWATH-MS data. **(A)** Schematic overview of the experimental design and SEC-SWATH-MS analysis workflow. HeLa cells were blocked in either interphase or mitosis. Samples were subject to mild cell lysis (1) followed by protein complex fractionation via SEC (2). Each of the consecutively sampled fractions was separately subjected to a bottom-up proteomic workflow including tryptic-digestion and MS data acquisition by SWATH-MS (3). The resulting peptide-level quantitative profiles along the SEC dimension were used for proteoform analysis with COPF (4), followed by optional protein complex assessment (5). **(B)** Histogram of proteoform scores for the cell cycle SEC-SWATH-MS dataset. A set of 115 proteins was predicted by the algorithm to contain multiple proteoform groups. **(C)** Assessment of the reported proteoform groups. **(D)** Uniprot keyword and sequence feature enrichment analysis of the predicted set of reported proteoform groups.

For the COPF analysis, we took the original peptide-level elution profiles from Heusel and Bludau et al.^28^ as a starting point. We imported the data into the R-CCprofiler framework for subsequent data preprocessing as follows. First, we annotated peptides with their start and end position in the canonical protein sequence. Second, peptides with overlapping start and end positions (e.g. peptides resulting from missed cleavages) were reduced to one representative peptide, selected based on the highest overall intensity across the dataset. Third, missing values were imputed for fractions with a valid value in both the preceding and following SEC fraction. Fourth, peptides detected in fewer than three consecutive fractions and peptides with zero variance along the SEC dimension were removed. Finally, only proteins with two or more remaining peptides were considered for further analysis by COPF.

The proteoform score distribution for the remaining 5451 proteins of the SEC-SWATH-MS dataset is shown in Figure 3B. We manually selected a conservative score threshold of 0.25 based on the criterion that the threshold precedes the exponential increase in the number of proteoforms observed at scores ≤ 0.2 (Supplementary Figure 1A and B). Based on this selected threshold, the algorithm predicted 115 proteins with functional proteoform groups, of which 86 (75%) are annotated with multiple isoforms in UniProt (Fisher’s exact test: odds ratio = 1.7, p-value = 0.006 when compared to the entire SEC-SWATH-MS dataset). A comparison with an independent study on phospho-signaling across cell cycle conditions^35^ showed that the 115 proteins are significantly enriched for cell-cycle regulated phosphosites (Fisher’s exact test: odds ratio = 7.3, p-value = 2×10-24, also see Supplementary Figure 1B). Proximity analysis of identified peptides within the protein sequence revealed that the proteoforms for 24 proteins (21%) were significantly closer in sequence proximity than expected by chance (p-value ≤ 10%) and proteoforms for an additional 43 proteins (37%) scored among the lowest 10% of possible p-values, given the number of peptides in the protein (for details also see Method section). We further analyzed the dataset with respect to proteoforms that associate with different complexes (assembly-specific) and to proteoforms that differ between cell-cycle states (cell cycle specific). Proteoform groups for 41 proteins (36%) could be classified as assembly-specific, since we observed them in multiple distinct assembly states resolved along the SEC dimension (for details see Method section). Additionally, COPF predicted proteoform groups of 53 proteins (46%) that were significantly differentially expressed between the two cell cycle stages (log2-fold change ≥ 1 and Benjamini-Hochberg (BH) adj. p-value ≤ 0.05). A summary of the proteoform characterization is provided in Figure 3C.

As a final assessment of the global characteristics of the set of proteins detected as having multiple proteoform groups, we performed an enrichment analysis (Figure 3D). We observed that the protein set is highly enriched in phosphoproteins. We also detected a significant enrichment in proline and poly-serine sequence features. This is particularly interesting because many cyclin dependent kinases act on sequence motifs including proline and serine^36–40^. Additionally, the protein set was enriched for proteins related to rearrangement of the cytoskeleton. This result is in agreement with cellular rearrangements required during mitosis. Finally, an enrichment of proteins with UniProt annotated sequence-(e.g. known polymorphisms) and splice-variants is observed.

In addition to these global insights, our dataset provided a rich source of new biological information. In the following we present selected examples that highlight different mechanisms generating proteoforms as well as different functional associations of proteoforms with either the cell cycle or protein assembly.

The first example is the proteasome subunit beta type-7 (PSMB7, UniProt ID: Q99436) which is convincingly resolved in our SEC data. The proteasome assembly line is a well-studied yet still heavily investigated system. Figure 4A shows a simplified schematic of the process. A key step in the prevalent model of 20S particle assembly is the integration of PSMB7 as the last beta subunit, triggering proteolytic cleavage of its pro-peptide, followed by formation of the full 20S core proteasome complex^41^. Our data-driven COPF strategy identified two assembly specific proteoforms for PSMB7 (also see Supplementary Figure 2A). The SEC profiles of all detected PSMB7 peptides are shown in Figure 4B. The peptides assigned to the two different proteoform groups are highlighted in blue and orange. While the peptides of both proteoform groups participate in the lower molecular weight peak around fraction 3 3, only peptides of the orange proteoform group were detected in the higher molecular weight peak around fraction 25. From protein co-elution analysis (Supplementary Figure 2C) as well as our previous study^21^, we know that the peak group around fraction 33 corresponds to a proteasome assembly intermediate and the peak group around fraction 25 corresponds to the full 20S core proteasome. Checking the location of the detected peptides along the PSMB7 sequence reveals that the peptides of the blue proteoform correspond to the two N-terminal peptides (Figure 4C, also see Supplementary Figure 2B). The second peptide (TGTTIAGVVYK) spans the known proteolytic cleavage site of the PSMB7 pro-peptide. To verify our finding, we performed a targeted re-extraction of the semi-tryptic peptide (TTIAGVVYK) that is produced by tryptic digestion of the processed, short proteoform using Skyline^42,43^. The extracted signal of this semi-tryptic peptide is highlighted in green in Figure 4B and C (also see Supplementary Figure 3). In contrast to the fully tryptic (blue) peptide, the cleaved peptide sequence co-elutes with the peptides of the orange proteoform group, indicating that the processed form is integrated in the 20S proteasome core complex as expected and identifying the precise location of the proteolytic processing that generates the proteoform.

**Figure 4.**
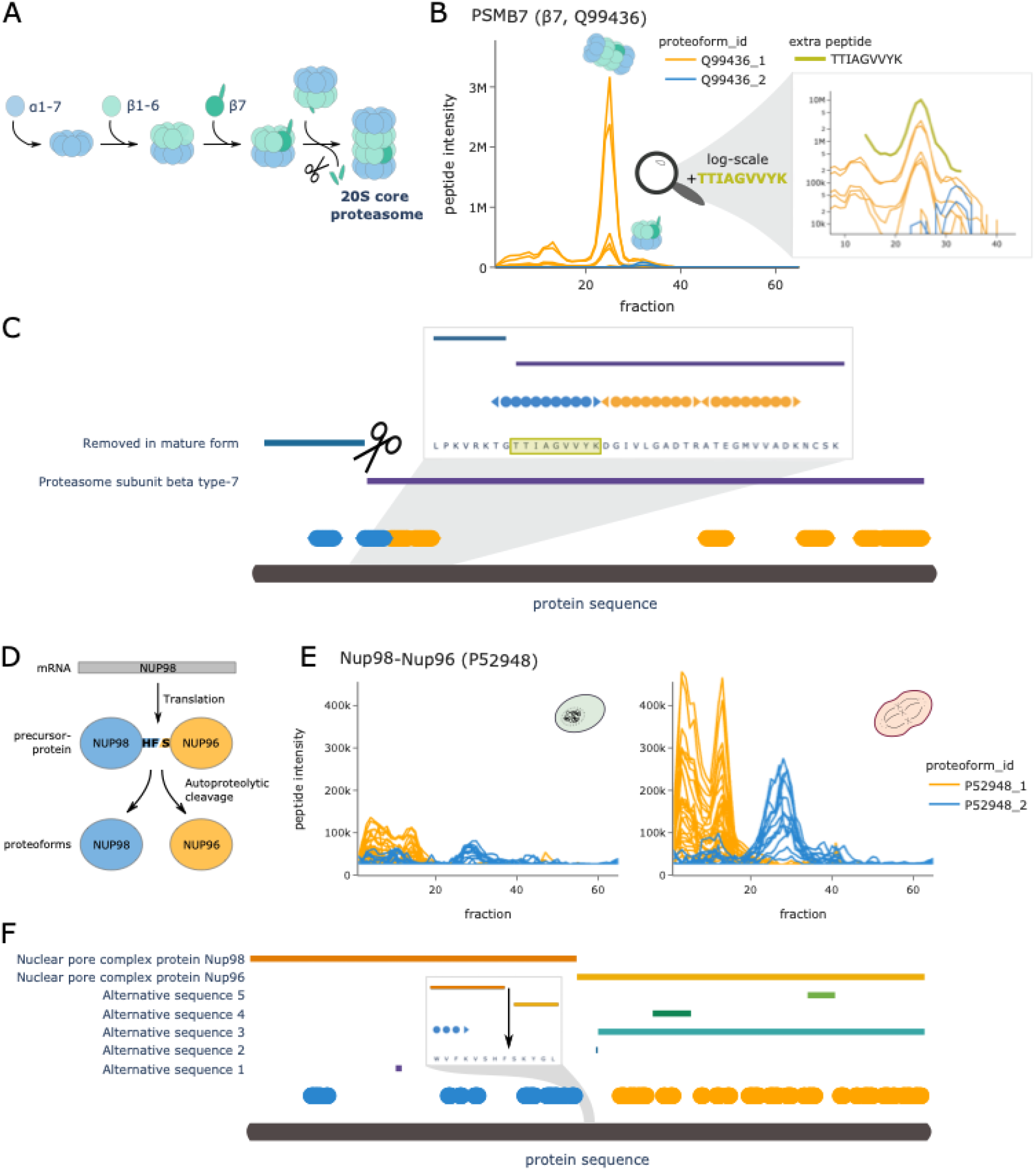
COPF results PSMB7 and NUP98/96. **(A)** Schematic overview of the proteasome assembly line. **(B)** Peptide profiles of the proteasome subunit beta type-7 (ß7, PSMB7, UniProt ID: Q99436) in interphase. Peptides of the two assigned proteoform groups are colored in orange and blue. A zoomin on fraction 10 to 40 is shown in log-scale, including an additional semi-tryptic peptide TTIAGVVYK that represents the N-terminal tryptic peptide of the processed proteoform (green). Please note that the abundance values between the orange and blue peptides cannot be directly compared to the semi-tryptic peptide TTIAGVVYK in green because of the separate analysis platforms used. **(C)** Protein sequence plot for PSMB7. Sequence coverage and position of the detected peptides of the two assigned proteoform groups are indicated in blue and orange. Pro-peptide and chain information from uniport are indicated as horizontal bars. The know pro-peptide cleavage site is indicated by scissors. A zoom-in representation is provided for the region around the annotated cleavage site. The semi-tryptic peptide from panel b is highlighted in green. **(D)** Schematic overview of the NUP98 and NUP96 proteoform biogenesis from the same mRNA and precursor protein (UniProt ID: P52948). **(E)** Peptide profiles of the NUP98 gene product in interphase (left) and mitosis (right). Peptides of the two assigned proteoform groups are colored in orange and blue. **(E)** Protein sequence plot for the NUP98 gene product. Sequence coverage and position of the detected peptides of the two assigned proteoform groups are indicated in blue and orange. Chain information and alternative sequence isoforms from uniport are indicated as horizontal bars. The known autocatalytic cleavage site of NUP98 is shown as an arrow in a zoom-in on the relevant sequence region.

The nuclear pore complex (NPC) protein Nup98-Nup96 (UniProt ID: P52948) presents a second example of the capacity of COPF for data-driven proteoform assignment. The Nup98-Nup96 proteoform groups were identified by the algorithm as both assembly and cell cycle specific (Figure 4D). The Nup98 gene is known to encode a 186 kDa precursor protein that undergoes autoproteolytic cleavage, to generate a 98 kDa nucleoporin (NUP98) and a 96 kDa nucleoporin (NUP96) (Figure 4D)^44–47^. While NUP96 is an important scaffold component of the NPC, NUP98 has diverse functional roles during mitosis. Previously, we showed the up-regulation of the Nup107-160 sub-complex (Corum ID: 87) in mitosis as compared to interphase (Figure 6H in^28^). There, we stated that the protein product of the Nup98 gene is up-regulated together with the other members of the Nup107-160 sub complex, i.e. proteins SEC13 (P55735), NUP107 (P57740), NUP160 (Q12769), NUP43 (Q8NFH3), NUP37 (Q8NFH4), NUP133 (Q8WUM0) SEH1L (Q96EE3) and NUP85 (Q9BW27). Here our purely data-driven approach was capable of correctly grouping peptides according to the two known proteoforms NUP98 and NUP96 (Figure 4E and F, also see Supplementary Figure 2D and E). Based on this assignment we can now also demonstrate that only the NUP96 proteoform integrates into the Nup107-160 sub-complex and follows its behavior across the cell cycle (Supplementary Figure 2F). This is in line with reports from previous studies^48^.

In addition to the above-mentioned examples where proteoforms were derived from enzymatic or autoproteolytic cleavage, the COPF algorithm could further detect proteoforms derived from alternative splicing, exemplified by the nuclear autoantigenic sperm protein (NASP, UniProt ID: P49321). Previous studies reported two alternative-splicing derived proteoforms of the NASP protein that can be detected in transformed cell lines: a somatic form (sNASP) and a shorter, testicular form (tNASP)^49^. In our HeLa cell cycle dataset COPF detected two assembly-specific proteoform groups that correctly matched the annotated proteoforms sNASP and tNASP (Figure 5A, also see Supplementary Figure 4A). The two distinct peptide peak groups in different molecular weight regions of the SEC separation indeed confirm previous findings that the two proteoforms engage in different assemblies.

**Figure 5.**
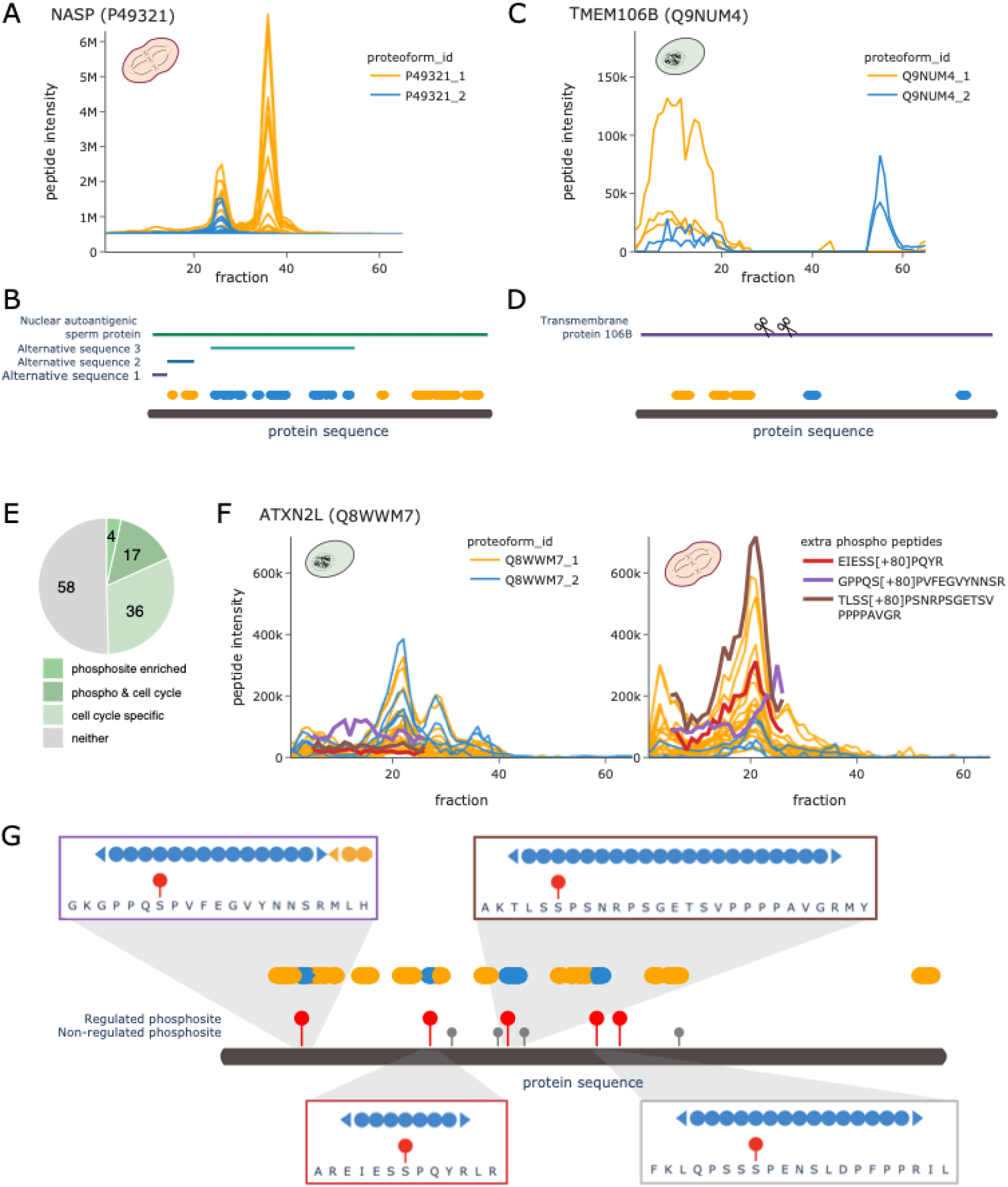
COPF results for NASP, TMEM106B and ATXN2L. **(A)** Peptide profiles of the nuclear autoantigenic sperm protein (NASP, UniProt ID: P49321) in mitosis. Peptides of the two assigned proteoform groups are colored in orange and blue. **(B)** Protein sequence plot for NASP. Sequence coverage and position of the detected peptides of the two assigned proteoform groups are indicated in blue and orange. Chain information and alternative sequence isoforms from uniport are indicated as horizontal bars. **(C)** Peptide profiles of the transmembrane protein 106B (TMEM106B, UniProt ID: Q9NUM4) in interphase. Peptides of the two assigned proteoform groups are colored in orange and blue. **(D)** Protein sequence plot for TMEM106B. Sequence coverage and position of the detected peptides of the two assigned proteoform groups are indicated in blue and orange. Chain information from uniport is indicated as horizontal bar. Suggested catalytic cleavage sites^50,51^ are indicated by scissors. **(E)** Pie chart illustrating the number of proteins with a proteoform enriched in cell-cycle regulated phosphosites^35^ and their overlap with cell cycle specificity of the proteoform. **(F)** Peptide profiles of the Ataxin-2-like protein (ATXN2L, UniProt ID: Q8WWM7) in interphase (left) and in mitosis (right). Peptides of the two assigned proteoform groups are colored in orange and blue. Additionally, traces of subsequently extracted phosphopeptides are highlighted in red, violet and brown. Please note that the abundance values between the orange and blue peptides cannot be directly compared to the phosphopeptides because of the separate analysis platforms used. **(G)** Protein sequence plot for ATXN2L. Sequence coverage and position of the detected peptides of the two assigned proteoform groups are indicated in blue and orange. Cell-cycle regulated phosophosites are indicated by red sticks, while non-regulated phosphosites are indicated in grey^35^. Regions of individual phosphopeptides are visualized in separate zoom-in windows, matching the colors of the respective peptides in panel F.

Whereas the examples described above refer to well annotated proteoforms that we were able to resolve without including prior knowledge in the analysis, our findings also uncovered less-well understood proteoforms. Figure 5C and D show the profile and sequence location for peptides of Transmembrane protein 106B (TMEM106B, UniProt ID: Q9NUM4), which has no annotated sequence variants or specific post-processing steps annotated in UniProt. Nevertheless, COPF identified two clearly distinguishable, assembly-specific proteoforms (Supplementary Figure 4B). In recent literature, we found evidence that TMEM106B is a lysosomal membrane protein that, upon membrane integration, undergoes evolutionarily-conserved regulated intramembrane proteolysis^50,51^. This involves a two-step mechanism where the luminal domain is first cleaved off by an unknown enzyme at amino-acid (AA) residue 127, followed by a second cleavage (at AA 106) of the N-terminal fragment that is still anchored in the membrane (cleavage sites are indicated by the scissors in Figure 5D). Our data suggest that the peptides of the blue proteoform group belong to the C-terminal luminal domain, eluting separately in the low molecular weight range, likely consistent with a monomeric form (expected monomeric MW = 31 kDa, observed elution at ~ 42 ± 10 kDa). The high molecular weight signal likely corresponds to the full protein integrated into the lysosomal membrane. Enrichment analysis of co-eluting proteins in the high molecular weight range shows an enrichment in membrane-associated proteins (Supplementary Figures 4C and D).

The strong enrichment of phospho-proteins in general and more specifically of proteins with previously reported cell cycle regulated phosphosites^35^ in the set of proteins with proteoform groups was a significant finding in the global analysis of COPF results (Figure 3D and Supplementary Figure 1B). To follow-up on these results, we set out to identify the specific proteoform groups that are enriched in cell cycle regulated phosphosites. For this, we performed enrichment analyses in each of our predicted proteoform groups to test if they are enriched for cell cycle regulated phosphosites as determined by Karayel et al.^35^. In total, 21 out of the 115 proteins with multiple proteoform groups (18%) had one proteofrom group significantly enriched in previously annotated cell cycle regulated phosphosites, thus suggesting that these proteoforms might be phosphorylation-derived. 17 (81%) of these proteins also turned out to be cell cycle stage specific (log2-fold change ≥ 1 and Benjamini-Hochberg (BH) adj. p-value ≤ 0.05, Figure 5E). One such example is the Ataxin-2-like protein (ATXN2L, UniProt ID: Q8WWM7). While peptides of the first proteoform group (orange) are higher-abundant in mitosis (log2-fold change = 1.38 and BH adj. p-value = 0.002), the second proteoform (blue) is significantly lower-abundant(log2-fold change up to −2.66 and Benjamini-Hochberg (BH) adj. p-value = 0.005) (Figure 5F). Four out of five regulated phosphosites^35^ (Figure 5G) fell into sequence regions covered in our dataset, exactly matching the four peptides of the second proteoform (blue). Making use of the possibility to re-extract data from SWATH-MS maps, we performed a targeted analysis in Skyline to detect and quantify the expected phosphopeptides in our dataset (Supplementary Figure 6). As predicted, we could confirm mitosis-specific phosphorylation for two of the four peptides (EIESS[+80]PQYR and TLSS[+80]PSNRPSGETSVPPPPAVGR, Figure 5F). One phosphopeptide (GPPQS[+80]PVFEGVYNNSR) had only a weak signal (purple) and the fourth could not be detected. Nevertheless, it is remarkable that the phosphopeptides could be detected and quantified in the samples given the long processing protocol without enrichment and without specific phosphatase inhibition treatment. These findings highlight that the COPF approach is capable of determining phospho-specific proteoform groups, given that at least two peptides are involved. Here, it is important to emphasize again that the strategy by which phospho-specific proteoform groups were detected with COPF directly links these detections to their biological relevance in the cell cycle.

### Tissue-specific proteoforms in SWATH-MS data of different mouse tissues

In comparison to the SEC-SWATH-MS dataset where native protein complexes are separated and analyzed in consecutive fractions, bottom-up proteomic datasets of unfractionated samples are more commonly available. Using a previously published SWATH-MS dataset of five different mouse tissues from eight BXD mice each^30^ (Figure 6A) we tested the assumption that the COPF strategy to identify functional proteoforms was also applicable to peptide intensity vs. sample data matrices from sample sets of sufficient size and variability between samples.

**Figure 6.**
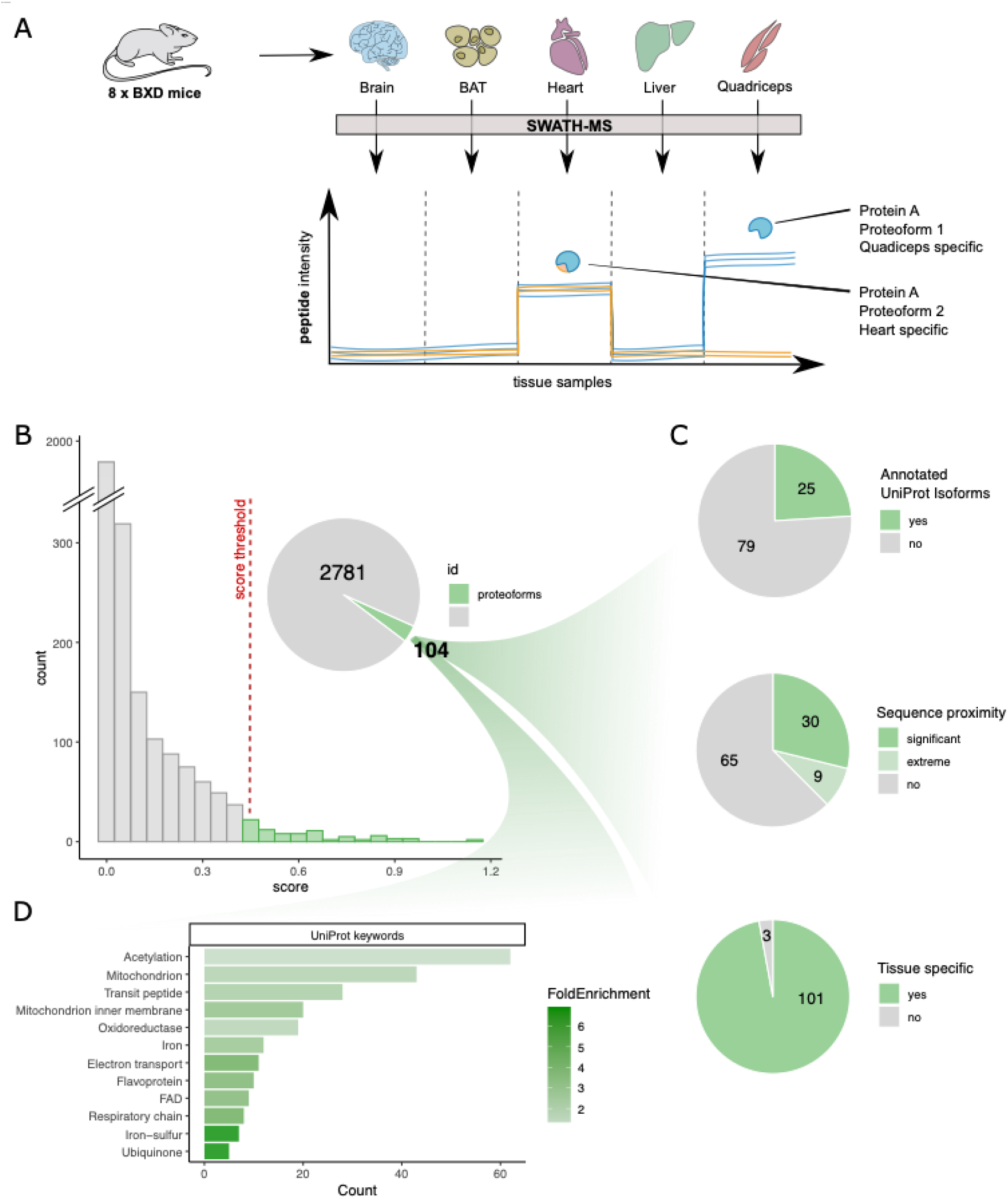
Global insights into tissue-specific proteoforms in full-proteome SWATH-MS data. **(A)** Schematic overview of the experimental design and analysis concept. Tissue samples were obtained from eight different strains of the BXD mouse genetic reference panel. The selected tissues were brain, brown adipose tissue (BAT), heart, liver and quadriceps. Each sample was separately processed by bottom-up proteomics using SWATH-MS. The resulting peptide-level quantitative profiles can be used to identify tissue-specific proteoform expression, here exemplified by muscle specific expression of a protein with a heart specific proteoform (blue and orange) and a quadriceps specific proteoform (only blue). **(B)** Histogram of proteoform scores for the mouse tissue SWATH-MS dataset generated by the COPF algorithm. 104 proteins were predicted with multiple proteoform groups. **(C)** Assessment of the reported proteoform groups. **(D)** Uniprot keyword enrichment analysis of the 104 proteins with multiple reported proteoform groups.

We initially imported the peptide-level data matrix (intensity vs. sample) into the R-CCprofiler framework and applied the same data processing steps as for the application of COPF for SEC-SWATH-MS data described above, except of the consecutive identification filter and missing value imputation. Proteins with two or more remaining peptides were considered for further analysis by COPF and resulted in the proteoform score distribution for 2885 proteins shown in Figure 6B. We manually selected a score threshold of 0.4 based on the criteria that the threshold precedes the exponential increase in the number of proteoforms observed for scores < 0.4 (Supplementary Figure 1C). Based on the selected threshold, the algorithm predicted 104 proteins with potential functional proteoform groups. 25 (24%) of these are annotated with multiple isoforms in UniProt. Peptide proximity analysis within the coding sequence further revealed that the proteoforms for 30 proteins (29%) were significantly closer in sequence proximity than expected by chance (p-value ≤ 10%) and proteoforms for an additional 9 proteins (9%) scored among the lowest 10% of possible p-values, given the number of peptides identified for the respective protein (for details also see Method section). Further evaluation of the detected proteoform groups revealed that proteoform groups for 101 proteins (97%) could be classified as tissue specific. This classification is based on the protein being differentially regulated when using tissue and predicted proteoform group information as prior knowledge for an ANOVA analysis (Bonferroni corrected p-value ≤ 0.01). A summary of the proteoform characterization is provided in Figure 6C.

We further performed an enrichment analysis of the proteins annotated with multiple proteoforms, showing that they are significantly enriched in keywords related to energy metabolism in mitochondria. This is in line with the expectation that energy metabolism is regulated differently between different tissues. Interestingly, many proteins are associated with acetylation.

Among the 104 proteins assigned to have functional proteoform groups, multiple biologically interesting instances stood out, exemplified by LIM domain-binding protein 3 (Ldb3, also known as Cypher, UniProt ID: Q9JKS4) and sorbin and SH3 domain-containing protein 2 (Sorbs2, UniProt ID: Q3UTJ2), respectively.

The protein Ldb3 was previously described as muscle specific^52^ and accordingly was found to be highly expressed only in heart and quadriceps samples of our dataset (Figure 7A). The COPF strategy clearly assigned the peptides of Ldb3 into two tissue-specific proteoform groups, indicated in orange and blue (Figure 7B). These proteoforms directly match the previously annotated splice variants of Ldb3, where peptides of the first proteofrom group, Q9JKS4-1, exactly map to the canonical sequence region that also has an alternative sequence variant (alternative sequence 1 in Figure 7C). To further validate this finding, we performed a targeted extraction of peptides in the alternative sequence region (peptides: VVANSPANADYQER and FNPSVLK) and could confirm their expression in quadriceps tissue (Figure 7B and C). These findings are in line with previous studies that reported tissue-specific expression of the alternative splice variant in skeletal muscle^52^.

**Figure 7.**
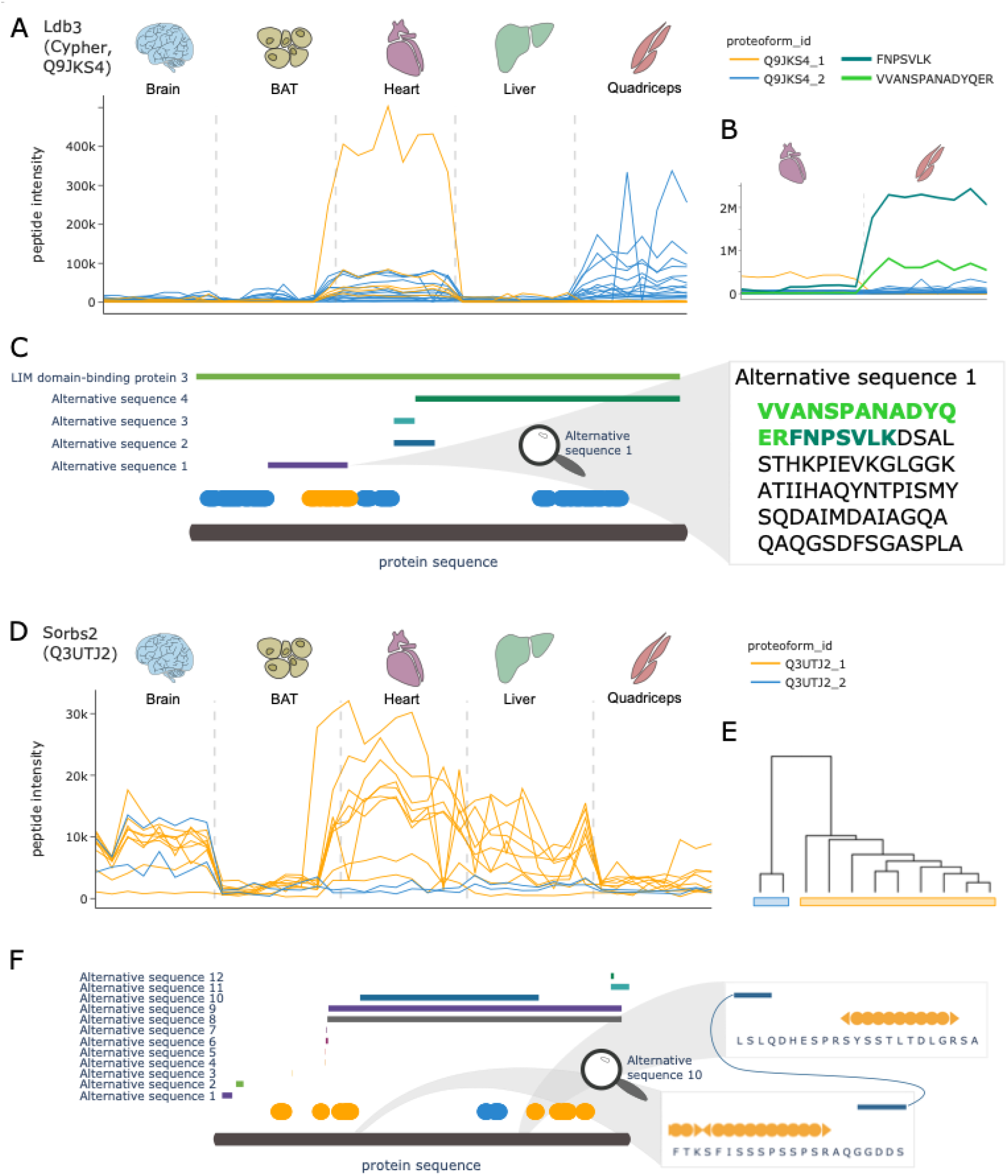
COPF results for Ldb3 and Sorbs2. **(A)** Peptide profiles of the LIM domain-binding protein 3 (Ldb3, also known as Cypher, UniProt ID: Q9JKS4). Peptides of the two assigned proteoform groups are colored in orange and blue. **(B)** Zoom-in on muscle tissue, specifically highlighting two additionally extracted peptides VVANSPANADYQER (light green) and FNPSCLK (dark green) that are specific to the known skeletal muscle-specific splice isoform of Ldb3. Please note that the abundance values between the orange and blue peptides cannot be directly compared to the additionally extracted peptides in green because of the separate analysis platforms used. **(C)** Protein sequence plot for Ldb3. Sequence coverage and position of the detected peptides of the two assigned proteoform groups are indicated in blue and orange. Chain information and alternative sequence isoforms from uniport are indicated as horizontal bars. The zoom-in on alternative sequence 1 shows the sequence of the skeletal muscle-specific splice isoform. The two peptides from panel B are highlighted in light and dark green respectively. **(D)** Peptide profiles of the sorbin and SH3 domain-containing protein 2 (Sorbs2, UniProt ID: Q3UTJ2). Peptides of the two assigned proteoform groups are colored in orange and blue. **(E)** Clustering dendogram for Sorbs2. **(F)** Protein sequence plot for Sorbs2. Sequence coverage and position of the detected peptides of the two assigned proteoform groups are indicated in blue and orange. Alternative sequence isoforms from uniport are indicated as horizontal bars. A zoom-in on the terminal regions of alternative sequence 10 shows that the orange proteoform does not cover its sequence region.

For Sorbs2 (Figure 7D) our fully data-driven approach assigned the peptides to two clearly distinct proteoform groups (Figure 7E). While the peptides of the orange proteoform group are abundant in brain, heart and liver tissues, the peptides of the second, blue proteoform group are observed exclusively in the brain. Mapping the identified peptides to the canonical protein sequence and matching them with annotated sequence variants (Figure 7F), it can be observed that the two blue peptides map to the region of a brain-specific splice variant (alternative sequence 10) that includes an exon that has previously been shown to only be expressed in brain tissue and exclusively in neurons^53–55^.

### A platform for manual exploration of the predicted functional proteoform groups

The examples discussed above only represent a fraction of the wealth of biological information that can be extracted from the COPF analysis of either the SECSWATH-MS or the mouse tissue proteomic datasets. To enable researchers to further explore the results in greater depth and to gain an insight into the sensitivity of the approach, we provide an online platform for manual data exploration which is openly available at: http://proteoformviewer.ethz.ch/ (password only during revision: ***).

We further provide all functional capabilities of COPF in an extended version of our software framework CCprofiler available at: https://github.com/CCprofiler/CCprofiler.

## Discussion

Multiple mechanisms including alternative splicing, protein processing and modification and the association of proteins in functional protein complexes contribute to the enormous expansion of the molecular diversity expressed from the ~ 20,000 protein coding genes of the human genome, and by analogy the genomes of other eukaryotic species. To date, proteomics has primarily operated in a gene-centric mode by focusing on the identification and quantification of proteins that can be assigned to their coding genes. The systematic resolution of proteoforms derived from individual genes in terms of their structure and function has remained largely elusive.

Until recently, proteoforms have mainly been inferred from transcriptomics data of next-generation sequencing (NGS) studies that aim to discover different mRNA transcripts of the same gene, mostly derived from alternative splicing. In these studies, it is assumed that the alternative transcripts are further translated to protein sequences, which is not always the case^56,57^. One approach to study proteoforms directly on the protein level is to generate and apply proteoform-specific antibodies, which can only be achieved at fairly low throughput. One recent example represents a study that characterized alternative CD6 isoforms generated by alternative-splicing^58^. However, the systematic identification of proteoforms on a proteome wide scale has only recently been enabled by MS based technologies.

Impressive recent progress in top-down proteomics has led to the identification and characterization of a few thousand proteoforms in parallel. Studies reported the identification of more than 3,000 unique proteoforms originating from up to ~1,000 individual genes^13,14^. Gaining deeper proteoform coverage by top-down proteomics is challenged by the limitations of current separation techniques, the MS and MS/MS analysis of large ions, and the interpretation of the resulting spectra by available analysis software^12,15^. In addition to the technical limitations, it has so far been difficult to systematically evaluate the functional relevance of the detected proteoforms.

Because the connection between protein and peptide is lost at an early stage of bottom-up proteomics, this method has mostly been considered unsuitable for the analysis of proteoforms to date. In this study we present a novel analysis concept and software we term COPF that i) systematically assigns peptides identified by bottom-up proteomics to proteoform groups and ii) directly links their identification to the biological context of the dataset at hand. We define a proteoform group as a group of peptides, derived from the same gene, that co-vary across large multi-condition datasets. COPF follows a fully data-driven approach, which necessitates a change in relative abundance between proteoform groups across the studied samples in order to enable proteoform group detection. The approach therefore directly couples detection to the assessment of a biological context.

An important distinction between functional proteoform groups assigned by COPF and those determined by top-down proteomics approaches is that COPF does not fully characterize the proteoform’s complete primary amino acid sequence and all of its modifications. It merely determines whether peptides exist that can differentiate the different biological contexts of a protein. Importantly, proteoform groups detected by COPF can directly imply a functional consequence depending on the study design. The power of the method is thereby directly linked to specific dataset properties. SEC-SWATH-MS data, as presented in this study, for example provides the opportunity to link the detection of proteoform groups directly to protein complex assembly, a property which makes such datasets unique and especially interesting for systematic proteoform investigation by COPF. A second factor based on which COPF can detect proteoform groups and distinguish functional associations is the biological context of a study, namely from the different biological conditions at hand. In the two datasets presented herein, these correspond to cell cycle- and tissue-specific proteoform groups.

A key challenge during the development of COPF was the lack of a ground truth dataset with known functional proteoforms. To nevertheless enable a performance evaluation of our software, we generated different *in silico* reference datasets. Our analysis demonstrated that COPF can both identify mixed proteins and assign peptides to the correct origin (Figure 2A-C). Further simulation studies allowed us to investigate the different parameters that influence the sensitivity of our approach. We could demonstrate that reducing technical variation between samples increases the sensitivity of mixed protein detection. Increased technical variation could further be rescued by increasing the number of peptides in a specific proteoform group (Figure 2D-F). These findings suggest that a high sequence coverage per protein as well as high reproducibility and quantitative accuracy are the key determinants for a successful proteoform group detection in bottom-up datasets. In this study, COPF was applied to two datasets generated by SWATH-MS. When compared to more classical data-dependent acquisition (DDA), DIA (or SWATH-MS) has the advantage of both increased data completeness and quantitative accuracy^59^. Recent developments on MS instrument level as well as in data acquisition and analysis promise further improvements with regard to proteome and sequence coverage^60,61^. Based on the results of the simulation benchmark, it can be expected that COPF performance and sensitivity will improve in concert. In addition to these technical considerations about sequence coverage and quantitative accuracy, it is also important to consider expected effect sizes of proteoform differences that can be observed in a given dataset. Differential SEC-SWATH-MS data, as presented herein, provides a rich source of variation based on the two components of protein complex fractionation and different biological conditions studied, i.e. interphase and mitosis. In studies with a more classical design of full proteome samples measured across different conditions, it is important to keep in mind that large effect sizes are required at the current technical standards of MS data acquisition. For this reason, different tissue types, as presented in this study, provide a good example for data with high abundance variation. Other promising study designs could for example be linked to sub-cellular localization maps or similar systems with a high degree of biological diversity. Applications might also extend to less well-defined study designs by mining large proteomic datasets available from proteomic databases such as Pride^62^. However, the scope of possible applications and its limitations still remain to be defined.

COPF application to both a cell cycle resolved SEC-SWATH-MS and a mouse tissue specific SWATH-MS dataset demonstrated that COPF can detect and correctly assign peptides to proteoform groups that correspond to well annotated sequence variants. In total, we found 115 candidate proteins for multiple proteoforms in the SEC-SWATH-MS dataset and 104 candidates in the mouse tissue data. In this study, thresholds were selected based on manual inspection and biological interpretation of the resulting proteoform groups (Supplementary Figure 1). A more systematic approach to further derive statistics about the false discovery rate was not possible at this stage, given that the properties of the underlying distributions are not yet fully understood. The reason for this is that the observed score distributions are a mixture of proteoforms with different signal-to-noise ratios and different fractions of peptides contributing to the signal. Our simulation studies (Figure 2D-F) indicated that these factors have a profound impact on the threshold that would be required for reaching a specified target false-discovery rate (FDR). Future studies will focus on a more systematic assessment of these factors and how to statistically assess the resulting score distributions. At this point we provide an interactive website that includes all data and results of our analysis for manual exploration and evaluation at http://proteoformviewer.ethz.ch/ (password only during revision: ***).

A limitation of COPF in its current implementation is that proteins will only be split into maximally two proteoform groups. It is expected that some proteins might have more than two functionally relevant proteoforms which could, at this point, not be resolved. Future work will focus on strategies towards a more fine-grained separation of proteins into a variable number of proteoform groups. This process will strongly benefit from a higher sequence coverage, that could potentially be achieved by newest DIA technologies in the near future^60,61^.

The presented examples in this study include proteoform groups generated by proteolytic cleavage (Figure 4A-C, Figure 5C-D), autocatalytic cleavage (Figure 4 D-F), alternative splicing (Figure 5A-B, Figure 7A-F) and multiple phosphorylations (Figure 5F-G). These examples demonstrate that the proposed strategy is, in principle, agnostic to the different mechanisms by which proteoforms can be generated inside the cell. This is in stark contrast to most other approaches that commonly investigate a subset of mechanisms. On one hand, proteoforms originating from alternative splicing are most commonly studied by proteogenomic approaches that combine RNA-sequencing with proteomics^63,64^. Classical PTM studies, on the other hand, are more commonly based on specific enrichment protocols that enable an in depth analysis of specific modifications, such as phosphorylations or ubiquitinations, with a focus on peptidoforms rather than proteoforms^3^. While the COPF strategy can, in principle, detect all types of proteoforms, the current implementation is limited by a minimum peptide set of two. The main reason is that it is difficult to confidently distinguish single outlier peptides from true biological signals. Future work towards improving outlier versus signal differentiation will therefore further increase the sensitivity of the COPF strategy and the scope of different proteoform groups that can be detected.

Although most presented examples confirm well annotated proteoforms, our approach has the unique feature of enabling their systematic co-detection and to directly enable the assessment of their relevance in the studied system, for example classifying them as assembly-, cell cycle- or tissue-specific. With the constantly increasing number of large-scale datasets and repositories generated by bottom-up proteomics^65–68^, there is a wealth of data waiting to be mined for new biology. We envision that our proteoform analysis concept will contribute to a paradigm shift towards the development of computational methods that directly couple discovery to biological context in such datasets. Strategies with such a direct link will enable easier interpretation of results and selection of promising follow-up candidates.

## Contributions

- IB, MF, MH, GR, BC, HR, and RA conceptualized the project
- IB, MF, MH, and HR conceptualized the COPF algorithm
- IB, MF, and YC implemented the COPF algorithm
- IB and YC performed the benchmarking
- IB performed the data analysis
- CD performed the Skyline analysis
- IB and CD interpreted the data with contributions from all authors
- IB created the website
- PP, BC, HR, and RA supervised the study
- IB and RA wrote the manuscript with contributions from all authors

## Acknowledgements

The authors thank Natalie De Souza for reading the manuscript and for providing valuable feedback. We also thank Uwe Schmitt, Patrick Pedrioli, Pascal Kägi for their help with setting up the website.

The project was supported by the SystemsX.ch project PhosphoNetX PPM (to R.A.), the European Research Council (ERC-20140AdG 670821 to R.A.) and the Swiss National Science Foundation project 310030E-173572 awarded under the DACH mechanism to R.A.. I.B. was supported by a Swiss National Science Foundation Postdoc.Mobility fellowship (P400PB_191046). G.R. was supported by grants P2EZP3_175127 and P400PB_183933 from the Swiss National Science Foundation. B.C.C. was supported by a Swiss National Science Foundation Ambizione grant (PZ00P3_161435).

## Methods

### Benchmarking by in silico sensitivity analysis

Details on the *in silico* benchmark are directly provided in the results section. A script with the complete sensitivity analysis is available on GitHub (https://github.com/ibludau/ProteoformAnanlysis/blob/main/PerformanceEvaluation/proteoformBenchmark_paper.R).

### Benchmarking simulation study

Details on the simulation are directly provided in the results section. A script with the complete simulation analysis is available on GitHub (https://github.com/ibludau/ProteoformAnanlysis/blob/main/PerformanceEvaluation/simulation.R).

### COPF analysis of the cell cycle SEC-SWATH-MS dataset

The peptide-level data and annotation of the cell cycle SEC-SWATH-MS dataset, E1709051521_feature_alignment.tsv and HeLaCCL2_SEC_annotation_full.csv, were downloaded from the Pride repository: https://www.ebi.ac.uk/pride/archive/projects/PXD010288. The data was loaded in R and imported as traces object, using *importMultipleConditionsFromOpenSWATH.* The proteins were further annotated with general information from UniProt using *annotateTraces.* Molecular weight (MW) calibration of the SEC fractions was performed based on measured standard proteins and their MWs by *calibrateMW* and *annotateMolecularWeight*. Peptide positions within the canonical protein sequence were determined by *annotatePeptideSequences*. Peptides from the same protein with similar start or end position, e.g. generated by missed cleavages, were summarized into a single peptide based on the highest intensity across the dataset by *summarizeAlternativePeptideSequences*. Missing values were imputed for fractions with a valid value in both the preceding and following SEC fraction using *findMissingValues* and *imputeMissingVals*. Peptides detected in fewer than three consecutive fractions across all replicates were excluded from further analysis *(filterConsecutiveIdStretches).* Finally, peptides with zero variance were removed and only proteins with multiple remaining peptides were kept for downstream analysis (f*ilterSinglePeptideHits*).

For COPF analysis, replicates of each condition were first integrated by *integrateReplicates* followed by appending SEC profiles across conditions by *combineTracesMutiCond.* Subsequently, all pairwise Pearson correlations between sibling peptides were calculated by *calculateGeneCorrMatrices*. Hierachical clustering based on an average linkage was performed by *clusterPeptides*. The tree was cut into two clusters by *cutClustersInNreal*. Each cluster is required to contain at least two peptides. Peptides that would form a single peptide cluster were marked as outliers. Finally, the proteoform score was calculated by *calculateProteoformScore*. Proteoform groups were annotated across all conditions and replicates by *annotateTracesWithProteoforms.*

To evaluate the sequence proximity of the resulting clusters, the *evaluateProteoformLocation* function was applied.

Scripts for the SEC-SWATH-MS COPF analysis are available on GitHub (https://github.com/ibludau/ProteoformAnanlysis/tree/main/CellCycleHela).

### Differential proteoform group analysis of the cell cycle SEC-SWATH-MS dataset

As a first step of differential proteoform group analysis, protein feature finding was performed. For this, peptide traces were summed across all conditions and replicates by *integrateTraceIntensities.* Peak groups along the SEC dimension were determined by findProteinFeatures (corr_cutoff = 0.9, window_size = 7, collapse_method = “apex_only”, perturb_cutoff = “5%”, rt_height = 1, smoothing_length=7, useRandomDecoyModel = TRUE, quantLevel = “protein_id”). Subsequently, protein features were resolved on the proteoform group level using *resolveProteoformSpecificFeatures* (minProteoformIntensityRatio = 0.1). Features were scored and filtered for a 5% FDR *(scoreFeatures,* FDR = 0.05).

The differential analysis was performed as previously described by Heusel et al.^28^ with minor modifications. Intensity values per condition and replicated were first extracted by *extractFeatureVals* and *fillFeatureVals*. The differential analysis was subsequently performed by *testDifferentialExpression.* Tests on the peptide level were finally aggregated to the proteoform group level by *aggregatePeptideTestsToProteoform*. Results were filtered by for a median log-2 fold change ? 1 and a Benjamini-Hochberg adjusted p-value ≤ 0.05.

A script for the differential SEC-SWATH-MS analysis is included on GitHub (https://github.com/ibludau/ProteoformAnanlysis/blob/main/CellCycleHela/06_differentialProteinFeatures_paper.R).

### COPF analysis of the mouse SWATH-MS dataset

The peptide-level quantitative data from Williams et al.^30^ was downloaded and only the whole proteome samples were selected for the analysis with CCprofiler and COPF. The quantitative data were loaded in R and imported as traces object, using *importPCPdata*. The proteins were further annotated with general information from UniProt using *annotateTraces*. Peptide positions within the canonical protein sequence were determined by *annotatePeptideSequences*. Peptides from the same protein with similar start or end position, e.g. generated by missed cleavages, were summarized into a single peptide based on the highest intensity across the dataset by *summarizeAlternativePeptideSequences*. Peptides with zero variance were removed and only proteins with multiple remaining peptides were kept for downstream analysis (f*ilterSinglePeptideHits*). Due to its suspicious properties, the protein with UniProt accession A2ASS6 was removed.

For COPF analysis, all pairwise Pearson correlations between sibling peptides were calculated by *calculateGeneCorrMatrices*. Hierachical clustering based on an average linkage was performed by *clusterPeptides*. The tree was cut into two clusters by *cutClustersInNreal.* Each cluster is required to contain at least two peptides. Peptides that would form a single peptide cluster were marked as outliers. Finally, the proteoform score was calculated by *calculateProteoformScore*. Proteoform groups were finally annotated by *annotateTracesWithProteoforms*.

To evaluate the sequence proximity of the resulting clusters, the *evaluateProteoformLocation* function was applied.

A script for the mouse tissue data analysis is available on GitHub (https://github.com/ibludau/ProteoformAnanlysis/blob/main/MouseTissue/GetMouseTissueProteoforms_paper.R).

### ANOVA analysis of the mouse SWATH-MS dataset

For the ANOVA analysis, we selected proteoforms based on a score cutoff of 0.4. Outlier peptides from the clustering were removed prior to proteoform quantification, which was performed by *proteinQuantification* (quantLevel = “proteoform_id”, topN = 1000, keep_less = TRUE). ANOVA analysis for each protein was performed using the aov function of the R stats library, applying following function: log2-intensity ~ tissue * proteoform. Multiple testing across all proteins was performed using a Bonferroni correction (p.adjust of the R stats library).

### Enrichment analyses

All presented enrichment analyses were performed on the DAVID website^69,70^. Results were filtered for a p-value of 0.01, a minimum count of 5 and a minimum fold-enrichment of 1.2.

### Sequence proximity analysis

The sequence proximity analysis evaluates if peptides assigned to the same proteoform group are in closer relative sequence proximity than expected for random peptide grouping. To test this hypothesis, the peptides of each protein are ranked by their relative peptide position start site (i.e. the position of the first amino acid of the given peptide in the canonical combined protein sequence). Subsequently, the normalized standard deviation of the derived peptide position ranks is calculated by dividing the standard deviation of the observed rank vector by the standard deviation of a uniform distribution of equal length. To estimate a probability of whether the observed proximity score is more extreme than expected by chance, we randomly shuffle the peptide ranks 1000 times for each protein and calculate the according proximity score for each of the permutations. An empirical p-value for each proteoform group is subsequently derived by dividing the number of random permutations with a more extreme proximity score compared to the real observation by the proximity score of the real observation. For proteins with very few detected peptides, it is impossible to reach statistical significance by means of a classical p-value. For these cases we implemented a second pseudo p-value that does not necessitate values ‘smaller or equal’ (as for the classical p-value), but that only considers ‘smaller’ values. This means that at a pseudo p-value of 5%, maximally 5% of the random permutations score better than the real observation. This criterion covers extreme cases with few peptides, which might still be taken into consideration for follow-up investigations. One example for a protein with two proteoforms that do not reach statistical significance in the sequence proximity analysis, because only two peptides in the short proteoform are too few, is PSMB7 (Supplementary Figure 2B). You can appreciate that the true observed proximity score is as low as statistically possible, therefore still potentially presenting an interesting candidate, as shown in this example. We report these extreme cases by stating that the protein scored among the lowest 10% of possible p-values.

### Phospho-enrichment analysis

To test our hypothesis that some of the proteoform groups detected by the cell cycle SEC-SWATH-MS analysis might be muti-phosphorylation derived, we retrieved the cell cycle phospho-proteomic dataset from Karayel et al.^35^. The dataset was downloaded from the original publication Supplementary Table 4 and filtered for ‘Mitosis/Interphase’ == TRUE and an absolute ‘Log2_ratios Mitosis/Interphase’ > 0.5. We used the remaining regulated phosphosites to test if proteins with multiple proteoforms in our cell cycle SEC-SWATH-MS dataset are enriched for these sites by performing Fisher’s exact test (fisher.test of the R stats library). To further test for each proteoform if more peptides than randomly expected contain a regulated phosphosite (as detected by Karayel et al.^35^), Fisher’s exact tests were performed for each proteoform in context of their protein. Proteins with a p-value ≤ 0.2 were considered interesting candidates where the proteoform could be related to multiple cell-cycle dependent phosphorylation events.

### Targeted analysis of selected peptides in Skyline

Selected raw data of both the cell cycle SEC-SWATH-SM and mouse tissue SWATH-MS data was downloaded from their respective Pride repositories, https://www.ebi.ac.uk/pride/archive/projects/PXD010288 and https://www.ebi.ac.uk/pride/archive/projects/PXD005044. Targeted extraction of the selected peptides was performed using Skyline version 4.1. The acquisition method was set to DIA and the isolation scheme was matched to the original publication of the datasets. Complete y and b ion series without retention time filtering were extracted for all peptides of interest. The data was than manually inspected for matching peak groups and the fragment ions were filtered for coelution. The quantification was subsequently based on the total area under the identified peak groups.

### Website

Data preprocessing and visualization for the dashboard was performed using the python programming language. The following libraries were utilized for data processing: numpy, pandas, scipy, re and pyteomics^71,72^. Parameter selection, tables and plots were generated using libraries from the HoloViz family of tools including: panel, holoviews, param, bokeh, plotly and matplotlib. Information about protein domains was retrieved from UniProt (https://www.uniprot.org/, accessed 22.06.2020 for human and 12.07.2020 for mouse), including following categories: ‘Chain’, ‘Domain’, ‘Alternative sequence’, ‘Propeptide’, ‘Signal peptide’ and ‘Transit peptide’.

## Notes

### Competing Interest Statement

The authors have declared no competing interest.

## References

1. Collins, F. S., Lander, E. S., Rogers, J. & Waterson, R. H. Finishing the euchromatic sequence of the human genome. Nature 431, 931–945 (2004).

2. van Straalen, N. M. & Roelofs, D. An Introduction to Ecological Genomics. An Introduction to Ecological Genomics (Oxford University Press, 2013). doi:10.1093/acprof:oso/9780199594689.001.0001.

3. Bludau, I. & Aebersold, R. Proteomic and interactomic insights into the molecular basis of cell functional diversity. Nature Reviews Molecular Cell Biology vol. 21 327–340 (2020).

4. Baralle, F. E. & Giudice, J. Alternative splicing as a regulator of development and tissue identity. Nat. Rev. Mol. Cell Biol. 18, 437 (2017).

5. Smith, L. M. et al. Proteoform: a single term describing protein complexity. Nat. Methods 10, 186 (2013).

6. Aebersold, R. et al. How many human proteoforms are there? Nat. Chem. Biol. 14, 206–214 (2018).

7. Kelemen, O. et al. Function of alternative splicing. Gene 514, 1–30 (2013).

8. Costa, V., Aprile, M., Esposito, R. & Ciccodicola, A. RNA-Seq and human complex diseases: Recent accomplishments and future perspectives. Eur. J. Hum. Genet. 21, 134–142 (2013).

9. Pistoni, M., Ghigna, C. & Gabellini, D. Alternative splicing and muscular dystrophy. RNA Biol. 7, 441–452 (2010).

10. Aebersold, R. & Mann, M. Mass spectrometry-based proteomics. Nature 422, 198 (2003).

11. Aebersold, R. & Mann, M. Mass-spectrometric exploration of proteome structure and function. Nature 537, 347 (2016).

12. Schaffer, L. V. et al. Identification and Quantification of Proteoforms by Mass Spectrometry. Proteomics 19, 1800361 (2019).

13. Tran, J. C. et al. Mapping intact protein isoforms in discovery mode using top-down proteomics. Nature 480, 254 (2011).

14. Anderson, L. C. et al. Identification and Characterization of Human Proteoforms by Top-Down LC-21 Tesla FT-ICR Mass Spectrometry. J. Proteome Res. 16, 1087–1096 (2017).

15. Toby, T. K., Fornelli, L. & Kelleher, N. L. Progress in Top-Down Proteomics and the Analysis of Proteoforms. Annu. Rev. Anal. Chem. 9, 499–519 (2016).

16. Nesvizhskii, A. I. & Aebersold, R. Interpretation of shotgun proteomic data: The protein inference problem. Molecular and Cellular Proteomics vol. 4 1419–1440 (2005).

17. Gillet, L. C. et al. Targeted Data Extraction of the MS/MS Spectra Generated by Data-independent Acquisition: A New Concept for Consistent and Accurate Proteome Analysis. Mol. & Cell. Proteomics 11, O111.016717 (2012).

18. Bruderer, R. et al. Extending the Limits of Quantitative Proteome Profiling with Data-Independent Acquisition and Application to Acetaminophen Treated Three-Dimensional Liver Microtissues. Mol. Cell. Proteomics 14, 1400–1410 (2015).

19. Collins, B. C. et al. Multi-laboratory assessment of reproducibility, qualitative and quantitative performance of SWATH-mass spectrometry. Nat. Commun. 8, 291 (2017).

20. Rosenberger, G. et al. Inference and quantification of peptidoforms in large sample cohorts by SWATH-MS. Nat. Biotechnol. 35, 781 (2017).

21. Heusel, M. et al. Complex-centric proteome profiling by SEC-SWATH-MS. Mol. Syst. Biol. 15, e8438 (2019).

22. Zhang, B., Pirmoradian, M., Zubarev, R. & Kall, L. Covariation of peptide abundances accurately reflects protein concentration differences. Mol. Cell. Proteomics 16, 936–948 (2017).

23. Webb-Robertson, B. J. M. et al. Bayesian proteoform modeling improves protein quantification of global proteomic measurements. Mol. Cell. Proteomics 13, 3639–3646 (2014).

24. Lukasse, P. N. J. & America, A. H. P. Protein inference using peptide quantification patterns. J. Proteome Res. 13, 3191–3199 (2014).

25. Forshed, J. et al. Enhanced information output from shotgun proteomics data by protein quantification and peptide quality control (PQPQ). Mol. Cell. Proteomics 10, (2011).

26. Meyer, J. G. Detection of Discordant Peptide Quantities in Shotgun Proteomics Data by Peptide Correlation Analysis (PeCorA). doi:10.1101/2020.08.21.261818.

27. Bamberger, C. et al. Deducing the presence of proteins and proteoforms in quantitative proteomics. Nat. Commun. 9, (2018).

28. Heusel, M. et al. A Global Screen for Assembly State Changes of the Mitotic Proteome by SEC-SWATH-MS. Cell Syst. 10, 133–155.e6 (2020).

29. Bludau, I. et al. Complex-centric proteome profiling by SEC-SWATH-MS for the parallel detection of hundreds of protein complexes. Nat. Protoc. 15, 2341–2386 (2020).

30. Williams, E. G. et al. Quantifying and localizing the mitochondrial proteome across five tissues in a mouse population. Mol. Cell. Proteomics 17, 1766–1777 (2018).

31. Consortium, T. U. UniProt: a worldwide hub of protein knowledge. Nucleic Acids Res. 47, D506–D515 (2018).

32. Röst, H. L. et al. OpenSWATH enables automated, targeted analysis of data-independent acquisition MS data. Nat. Biotechnol. 32, 219 (2014).

33. Ting, Y. S. et al. Peptide-Centric Proteome Analysis: An Alternative Strategy for the Analysis of Tandem Mass Spectrometry Data. Mol. Cell. Proteomics 14, 2301–2307 (2015).

34. Rosenberger, G. et al. Statistical control of peptide and protein error rates in large-scale targeted data-independent acquisition analyses. Nat. Methods (2017) doi:10.1038/nmeth.4398.

35. Karayel, Ö. et al. Comparative phosphoproteomic analysis reveals signaling networks regulating monopolar and bipolar cytokinesis. Sci. Rep. 8, (2018).

36. Moreno, S. & Nurse, P. Substrates for p34cdc2: In vivo veritas? Cell vol. 61 549–551 (1990).

37. Nigg, E. A. Cellular substrates of p34cdc2 and its companion cyclin-dependent kinases. Trends Cell Biol. 3, 296–301 (1993).

38. Songyang, Z. et al. Use of an oriented peptide library to determine the optimal substrates of protein kinases. Curr. Biol. 4, 973–982 (1994).

39. Errico, A., Deshmukh, K., Tanaka, Y., Pozniakovsky, A. & Hunt, T. Identification of substrates for cyclin dependent kinases. Adv. Enzyme Regul. 50, 375–399 (2010).

40. Alexander, J. et al. Spatial exclusivity combined with positive and negative selection of phosphorylation motifs is the basis for context-dependent mitotic signaling. Sci. Signal. 4, ra42 (2011).

41. Gu, Z. C. & Enenkel, C. Proteasome assembly. Cellular and Molecular Life Sciences vol. 71 4729–4745 (2014).

42. MacLean, B. et al. Skyline: an open source document editor for creating and analyzing targeted proteomics experiments. Bioinformatics 26, 966–968 (2010).

43. Pino, L. K. et al. The Skyline ecosystem: Informatics for quantitative mass spectrometry proteomics. Mass Spectrometry Reviews vol. 39 229–244 (2020).

44. Enninga, J., Levy, D. E., Blobel, G. & Fontoura, B. M. A. Role of nucleoporin induction in releasing an mRNA nuclear export block. Science (80-.). 295, 1523–1525 (2002).

45. Hodel, A. E. et al. The three-dimensional structure of the autoproteolytic, nuclear pore-targeting domain of the human nucleoporin Nup98. Mol. Cell 10, 347–358 (2002).

46. Fontoura, B. M. A., Blobel, G. & Matunis, M. J. A conserved biogenesis pathway for nucleoporins: Proteolytic processing of a 186-kilodalton precursor generates Nup98 and the novel nucleoporin, Nup96. J. Cell Biol. 144, 1097–1112 (1999).

47. Rosenblum, J. S. & Blobel, G. Autoproteolysis in nucleoporin biogenesis. Proc. Natl. Acad. Sci. U. S. A. 96, 11370–11375 (1999).

48. Beck, M. & Hurt, E. The nuclear pore complex: Understanding its function through structural insight. Nature Reviews Molecular Cell Biology vol. 18 73–89 (2017).

49. Richardson, R. T. et al. Characterization of the histone H1-binding protein, NASP, as a cell cycle-regulated somatic protein. J. Biol. Chem. 275, 30378–30386 (2000).

50. Nicholson, A. M. & Rademakers, R. What we know about TMEM106B in neurodegeneration. Acta Neuropathologica vol. 132 639–651 (2016).

51. Brady, O. A., Zhou, X. & Hu, F. Regulated intramembrane proteolysis of the frontotemporal lobar degeneration risk factor, TMEM106B, by signal peptide peptidase-like 2a (SPPL2a). J. Biol. Chem. 289, 19670–19680 (2014).

52. Huang, C. et al. Characterization and in vivo functional analysis of splice variants of cypher. J. Biol. Chem. 278, 7360–7365 (2003).

53. Zhang, Q. et al. Impaired dendritic development and memory in sorbs2 knock-out mice. J. Neurosci. 36, 2247–2260 (2016).

54. Kawabe, H. et al. nArgBP2, a novel neural member of ponsin/ArgBP2/vinexin family that interacts with synapse-associated protein 90/postsynaptic density-95-associated protein (SAPAP). J. Biol. Chem. 274, 30914–30918 (1999).

55. Lee, S. E., Kim, J. A. & Chang, S. NArgBP2-SAPAP-SHANK, the core postsynaptic triad associated with psychiatric disorders. Exp. Mol. Med. 50, 1–9 (2018).

56. Tress, M. L., Abascal, F. & Valencia, A. Alternative Splicing May Not Be the Key to Proteome Complexity. Trends Biochem. Sci. 42, 98–110 (2017).

57. Wan, Y. & Larson, D. R. Splicing heterogeneity: separating signal from noise. Genome Biol. 19, 86 (2018).

58. Santos, R. F., Oliveira, L., Brown, M. H. & Carmo, A. M. Domain-specific <scp>CD>/scp> 6 monoclonal antibodies identify <scp>CD>/scp> 6 isoforms generated by alternative-splicing. Immunology 157, imm.13087 (2019).

59. Bruderer, R. et al. Optimization of Experimental Parameters in Data-Independent Mass Spectrometry Significantly Increases Depth and Reproducibility of Results. Mol. & Cell. Proteomics 16, 2296LP–2309 (2017).

60. Meier, F. et al. Parallel accumulation-serial fragmentation combined with data-independent acquisition (diaPASEF): Bottom-up proteomics with near optimal ion usage. bioRxiv 656207 (2019) doi:10.1101/656207.

61. Bekker-Jensen, D. B. et al. A compact quadrupole-orbitrap mass spectrometer with FAIMS interface improves proteome coverage in short LC gradients. Mol. Cell. Proteomics 19, 716–729 (2020).

62. Perez-Riverol, Y. et al. The PRIDE database and related tools and resources in 2019: Improving support for quantification data. Nucleic Acids Res. 47, D442–D450 (2019).

63. Nesvizhskii, A. I. Proteogenomics: concepts, applications and computational strategies. Nat. Methods 11, 1114 (2014).

64. Sheynkman, G. M., Shortreed, M. R., Cesnik, A. J. & Smith, L. M. Proteogenomics: Integrating Next-Generation Sequencing and Mass Spectrometry to Characterize Human Proteomic Variation. Annu. Rev. Anal. Chem. 9, 521–545 (2016).

65. Wilhelm, M. et al. Mass-spectrometry-based draft of the human proteome. Nature 509, 582 (2014).

66. Wang, D. et al. A deep proteome and transcriptome abundance atlas of 29 healthy human tissues. Mol. Syst. Biol. 15, (2019).

67. Doll, S. et al. Region and cell-type resolved quantitative proteomic map of the human heart. Nat. Commun. 8, (2017).

68. Kim, M.-S. et al. A draft map of the human proteome. Nature 509, 575 (2014).

69. Huang, D. W., Sherman, B. T. & Lempicki, R. A. Systematic and integrative analysis of large gene lists using DAVID bioinformatics resources. Nat. Protoc. 4, 44–57 (2009).

70. Huang, D. W., Sherman, B. T. & Lempicki, R. A. Bioinformatics enrichment tools: Paths toward the comprehensive functional analysis of large gene lists. Nucleic Acids Res. 37, 1–13 (2009).

71. Goloborodko, A. A., Levitsky, L. I., Ivanov, M. V. & Gorshkov, M. V. Pyteomics-A python framework for exploratory data analysis and rapid software prototyping in proteomics. J. Am. Soc. Mass Spectrom. 24, 301–304 (2013).

72. Levitsky, L. I., Klein, J. A., Ivanov, M. V. & Gorshkov, M. V. Pyteomics 4.0: Five Years of Development of a Python Proteomics Framework. J. Proteome Res. 18, 709–714 (2019).

