## Supplementary Tables and Figures for "Systematic detection of functional proteoform groups from bottom-up proteomic datasets"

**Supplementary Table 1: Glossary.**

| Term | Definition |
|---|---|
| Proteoform | A protein species with a unique combination of amino acid sequence and post-translational modifications. Multiple alternative proteoforms can originate from the same gene locus. |
| Functional proteoform group | A group of peptides, derived from the same gene, that co-vary across a large multi-condition dataset. A proteoform group can, but does not have to resemble a unique, specific proteoform. |
| Protein with functional proteoform groups | A protein with at least two groups of peptides that show different patterns across a large multi-condition dataset. |

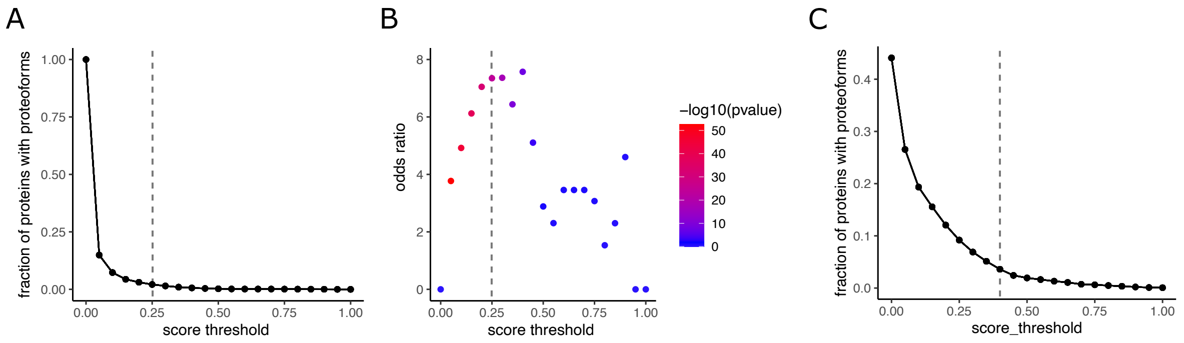


**Supplementary Figure 1. Criteria for the selection of proteoform score thresholds. (A)** Fraction of proteins with multiple proteoforms for different score thresholds in the cell cycle SEC-SWATH-MS dataset. The selected threshold of 0.25 is indicated as dashed vertical line. **(B)** Odds ratio of Fisher's exact test for the enrichment of regulated phosphosites within the set of proteins determined to have multiple proteoforms across different score thresholds. The selected threshold of 0.25 is indicated as dashed vertical line. **(C)** Fraction of proteins with multiple proteoforms for different score thresholds in the mouse tissue SWATH-MS dataset. The selected threshold of 0.4 is indicated as dashed vertical line.

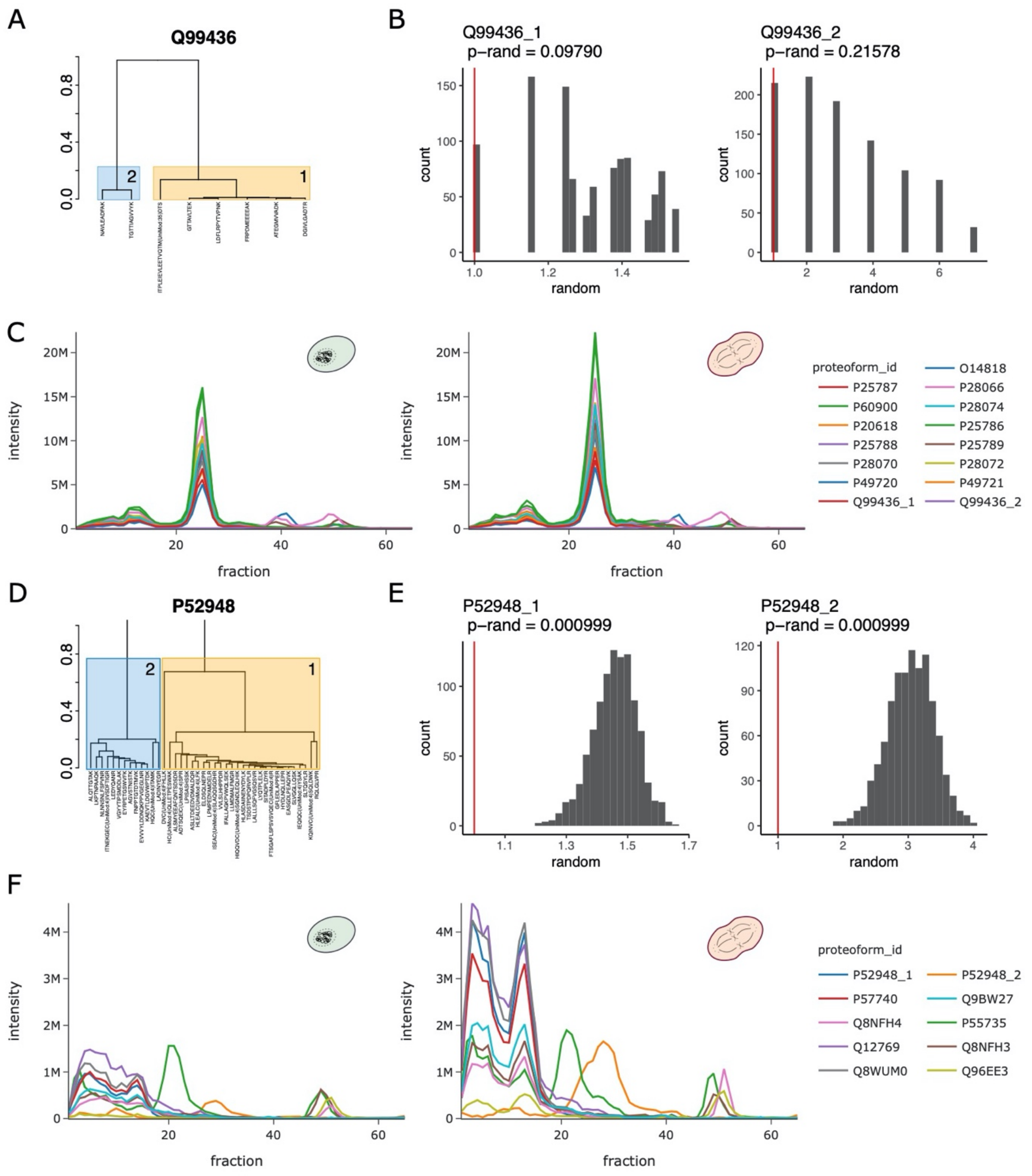


**Supplementary Figure 2. Additional information about COPF results for PSMB7 and NUP98/96. (A)** Clustering dendogram for proteasome subunit beta type-7 (β7, PSMB7, UniProt ID: Q99436). The two determined proteoforms are highlighted in orange and blue. **(B)** Histograms for the sequence proximity analysis of PSMB7 proteoforms 1 (left) and 2 (right). The grey bars illustrate the proximity score distribution for 1000-times randomly shuffled peptide ranks. The red vertical line indicates the score achieved on the true, observed peptide position rank. **(C)** Protein profiles for all annotated 20S proteasome subunits in interphase (left) and mitosis (right). **(D)** Clustering dendogram for the NUP98 gene product (UniProt ID: P52948). The two determined proteoforms are highlighted in orange and blue. **(E)** Histograms for the sequence proximity analysis of the NUP98 proteoforms 1 (left) and 2 (right). The grey bars illustrate the proximity score distribution for 1000-times randomly shuffled peptide ranks. The red vertical line indicates the score achieved on the true, observed peptide position rank. **(F)** Protein profiles for all annotated Nup107-160 sub-complex subunits in interphase (left) and mitosis (right).

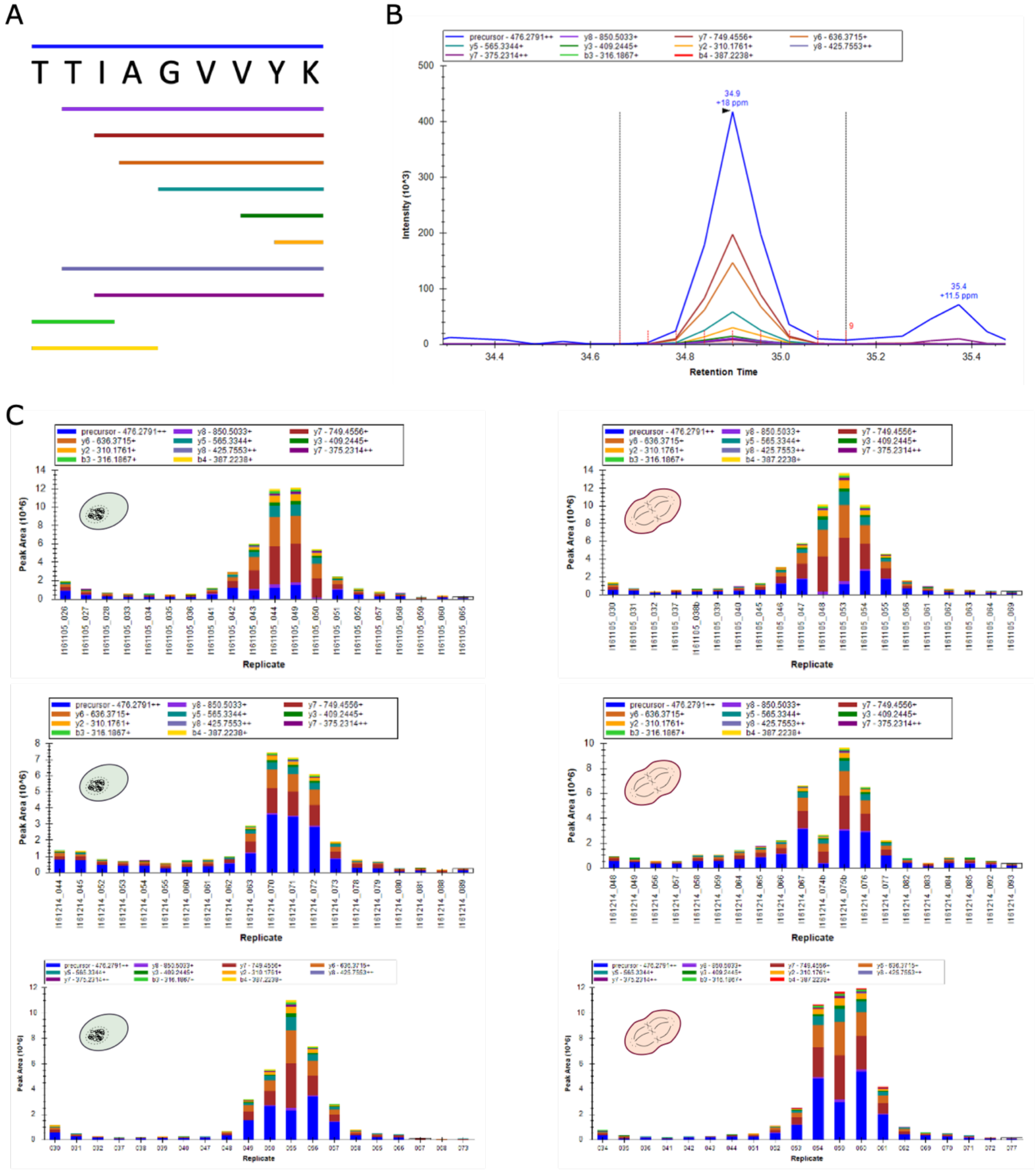


**Supplementary Figure 3. Skyline extraction of the semi-tryptic peptide generated by proteolytic cleavage of the pro-peptide of PSMB7. (A)** Peptide sequence of the expected semi-tryptic peptide (TTIAGVVYK) generated by proteolytic cleavage of the pro-peptide in PSMB7. The full precursor sequence is indicated in blue on top of the sequence while different fragment ions are indicated as horizontal lines below the sequence. **(B)** A representative fragment-ion peak group for TTIAGVVYK that was extracted in Skyline. **(C)** Quantitative bar plot of the extracted TTIAGVVYK signals across the three replicates in interphase (left) and mitosis (right).

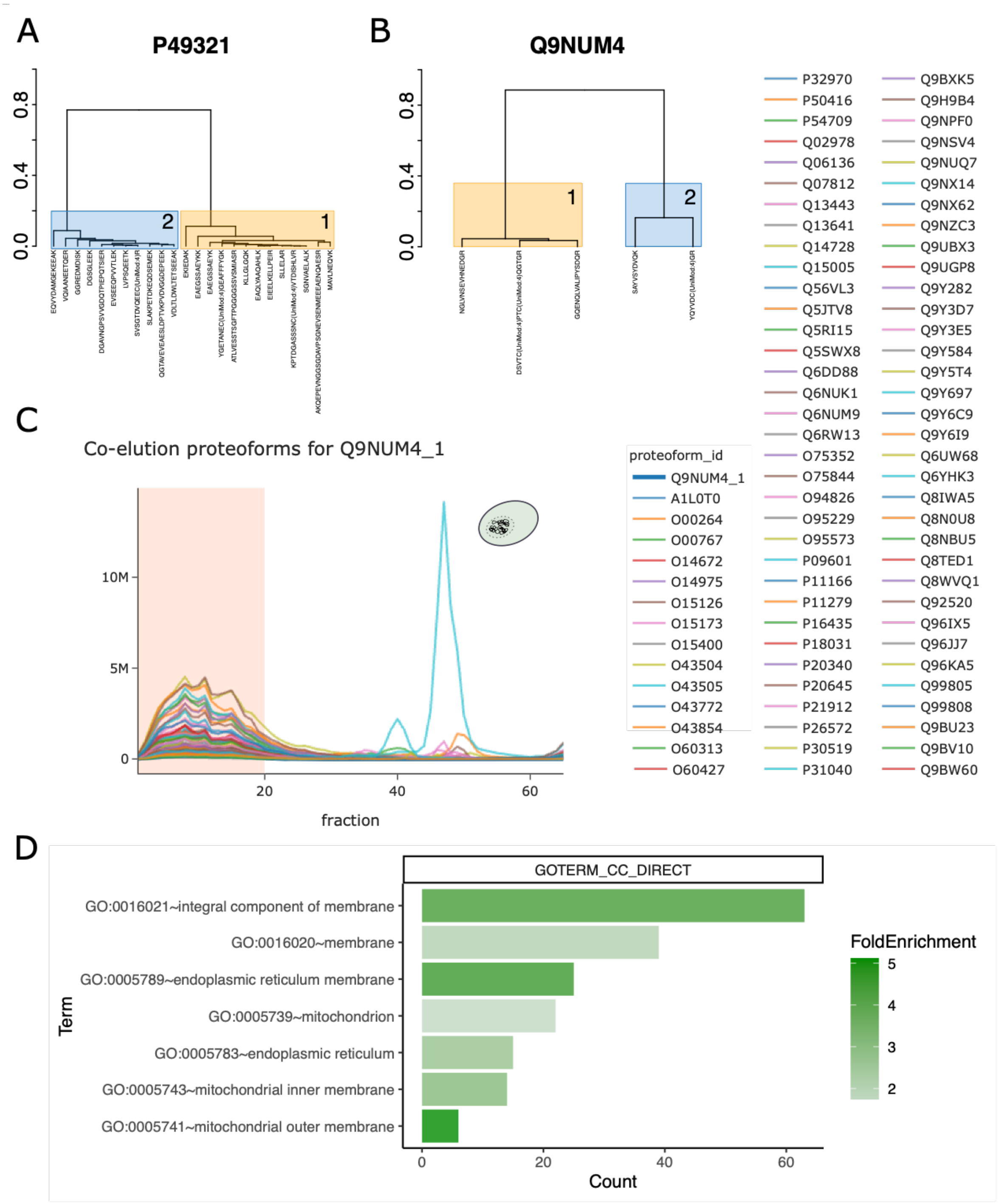


**Supplementary Figure 4. Additional information on proteoform groups detected for NASP and TMEM106B. (A)** Clustering dendogram for nuclear autoantigenic sperm protein (NASP, UniProt ID: P49321). The two determined proteoforms are highlighted in orange and blue. **(B)** Clustering dendogram for Transmembrane protein 106B (TMEM106B, UniProt ID: Q9UM4). The two determined proteoforms are highlighted in orange and blue. **(C)** High-correlating protein profiles for TMEM106B proteoform group 1 (Q9NUM4_1) in interphase. A fraction range from 1 to 20 and a minimum Pearson correlation of 0.95 were selected. **(D)** Enriched gene ontology (GO) based cellular components among the proteins that are highly correlated with TMEM106B proteoform group 1.

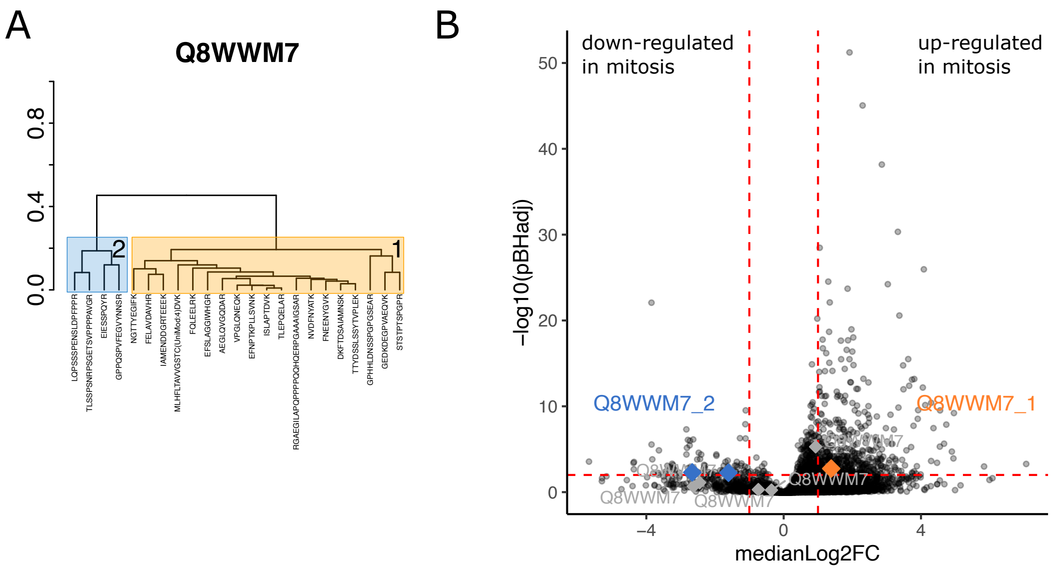


**Supplementary Figure 5. Additional information on proteoform groups detected for ATXN2L. (A)** Clustering dendogram for ataxin-2-like protein (ATXN2L, UniProt ID: Q8WWM7). The two determined proteoforms are highlighted in orange and blue. **(B)** Volcano plot showing the median log-2 fold change (medianLog2FC) over the negative Benjamini-Hochberg corrected p-value (-log10(pBHadj)). Each point represents a detected SEC elution peak for a specific proteoform group. The diamonds indicate the detected peaks for ATXN2L (UniProt ID: Q8WWM7). The colored diamonds are significantly up- or down-regulated signals (FDR ≤ 5% and absolute log-2 fold-change ≥ 1) for either of the two proteoform groups represented in orange and blue, respectively.

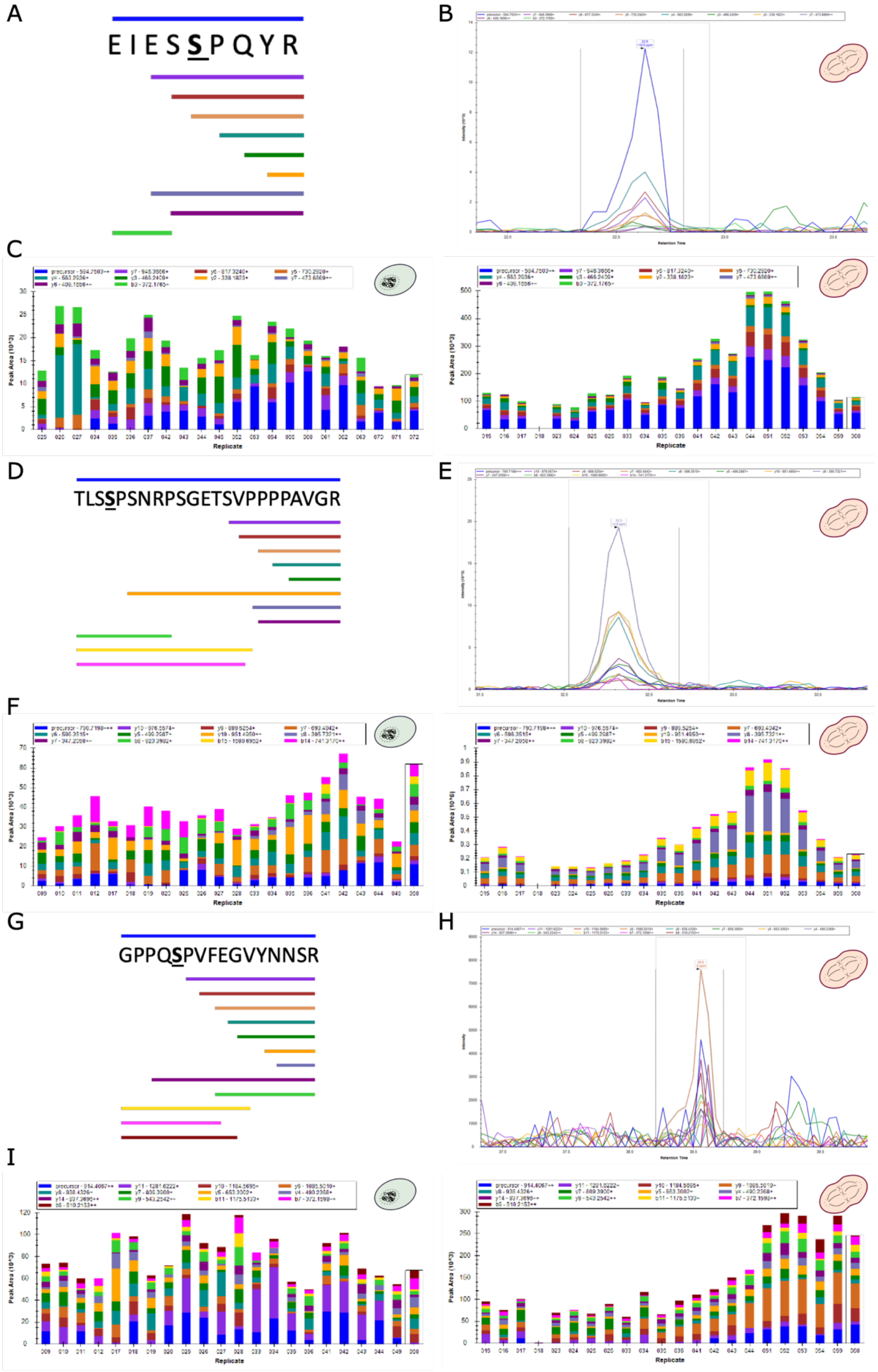
A
EIESSPQYR
B
C
Peak Area (10^3)
Replicate
D
TLSSPSNRPSGETSVPPPPAVGR
E
F
Peak Area (10^6)
G
GPPQSPVFEGVYNNSR
H
I

**Supplementary Figure 5. Skyline extraction of three phospho-peptides of ATXN2L. (A)** Peptide sequence of the expected phospho-peptide EIES**S**PQYR, the expected phospho-site is highlighted in bold. The full precursor sequence is indicated in blue on top of the sequence while different fragment ions are indicated as horizontal lines below the sequence. **(B)** A representative fragment-ion peak group for EIES**S**PQYR that was extracted in Skyline (mitosis sample). **(C)** Quantitative bar plots for the extracted TLS**S**PSNRPSGETSVPPPPAVGR signals across one representative replicate in interphase (left) and mitosis (right). **(D)** Peptide sequence of the expected phospho-peptide TLS**S**PSNRPSGETSVPPPPAVGR, the expected phospho-site is highlighted in bold. The full precursor sequence is indicated in blue on top of the sequence while different fragment ions are indicated as horizontal lines below the sequence. **(E)** A representative fragment-ion peak group for TLS**S**PSNRPSGETSVPPPPAVGR that was extracted in Skyline (mitosis sample). **(F)** Quantitative bar plots for the extracted TLS**S**PSNRPSGETSVPPPPAVGR signals across one representative replicate in interphase (left) and mitosis (right). **(G)** Peptide sequence of the expected phospho-peptide GPPQ**S**PVFEGVYNNSR, the expected phospho-site is highlighted in bold. The full precursor sequence is indicated in blue on top of the sequence while different fragment ions are indicated as horizontal lines below the sequence. **(H)** A representative fragment-ion peak group for GPPQ**S**PVFEGVYNNSR that was extracted in Skyline (mitosis sample). **(I)** Quantitative bar plots for the extracted GPPQ**S**PVFEGVYNNSR signals across one representative replicate in interphase (left) and mitosis (right).

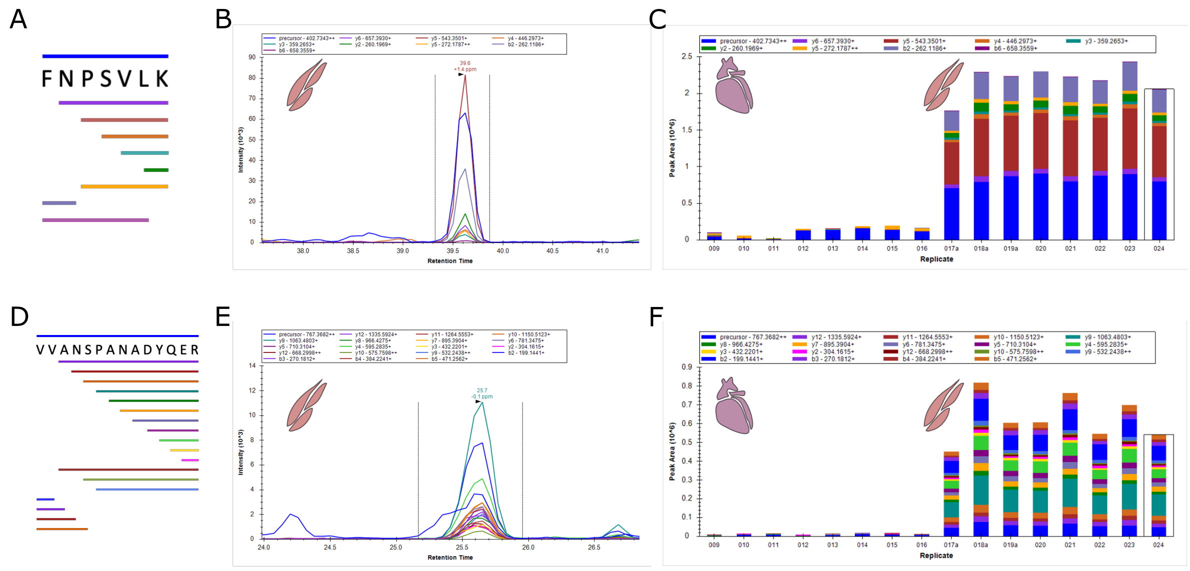


**Supplementary Figure 7. Skyline extraction of peptides specific to a known skeletal muscle-specific splice isoform of** Ldb3 (cypher)**. (A)** Peptide sequence of the expected skeletal muscle-specific splice isoform FNPSVLK. The full precursor sequence is indicated in blue on top of the sequence while different fragment ions are indicated as horizontal lines below the sequence. **(B)** A representative fragment-ion peak group for FNPSVLK that was extracted in Skyline from a quadriceps sample. **(C)** Quantitative bar plots for the extracted FNPSVLK signals across the heart (left) and quadriceps samples (right). **(D)** Peptide sequence of the expected skeletal muscle-specific splice isoform VVANSPANADYQER. The full precursor sequence is indicated in blue on top of the sequence while different fragment ions are indicated as horizontal lines below the sequence. **(E)** A representative fragment-ion peak group for VVANSPANADYQER that was extracted in Skyline from a quadriceps sample. **(F)** Quantitative bar plots for the extracted VVANSPANADYQER signals across the heart (left) and quadriceps samples (right).
